# *Entamoeba histolytica* Gal/GalNAc lectin intermediate subunit promotes inflammation and epithelial damage in intestinal amebiasis through its C3 region

**DOI:** 10.1101/2025.04.22.649943

**Authors:** Hongze Zhang, Shaokun Pan, Meng Feng, Dai Dong, Yanqing Zhao, Ruixue Zhou, Wenjie Li, Xunjia Cheng

**Affiliations:** Department of Medical Microbiology and Parasitology, School of Basic Medical Sciences, Fudan University, Shanghai, China; Shanghai Institute of Infectious Disease and Biosecurity, Fudan University, Shanghai, China

**Author notes:** Address correspondence to Xunjia Cheng,. Hongze Zhang, Shaokun Pan, and Meng Feng contributed equally to this work.

**Keywords:** *Entamoeba histolytica*, intestinal amebiasis, Gal/GalNAc lectin, single-cell RNA sequencing, macrophage, epithelial cell, inflammation, extracellular vesicle

## Abstract

*Entamoeba histolytica* is a common enteric protozoan that sometimes transitions from a symbiont to a pathogen, causing intestinal amebiasis and extraintestinal abscesses in the host. The early-to-intermediate stages of intestinal amebiasis are characterized by an intense inflammatory response, during which host innate immune defenses play an important role. Here, using the cecal infection model of C3H/HeNCrl mice, we performed single-cell RNA sequencing of cecal tissues and identified the prominent pro-inflammatory function of host macrophages in intestinal amebiasis. On the amoebic cell surface, Igl is an intermediate subunit of galactose- and N-acetyl-D-galactosamine-inhibitable lectins, which contributes to parasite adherence. We proposed that *E. histolytica* Igl diffuses along with amoebic extracellular vesicles and contacts host macrophages, binding to the TLR4 co-receptor MD2 through its C3 region and initiating the TLR4/MyD88/NF-κB inflammatory signaling pathway in host cells. Moreover, by inducing macrophage inflammation and cytokine production, *E. histolytica* indirectly impairs intestinal epithelium integrity through the Igl protein. To our knowledge, this is the first report showing a host single-cell atlas in the field of amebiasis research. In response to the parasite invasion during intestinal amebiasis, our study provides new insights elucidating the inflammatory formation and epithelial damage of gut mucosal immune system, and also conduces to the identification of potential drug targets for these amoebas.

## INTRODUCTION

Intestinal protozoan inhabitants play an important role in shaping the gut bacterial microbiota and maintaining host-microbe equilibrium in healthy individuals ^1, 2^. Protozoa of the genus *Entamoeba* colonize animals widely, and at least seven *Entamoeba* species are endobiotic in the human gut ^3^. Among them, the only one that can become virulent and cause serious damage to the host is *Entamoeba histolytica*, the causative agent of various amebiases such as amoebic colitis and liver abscesses ^4^. As a common intestinal protozoan prevalent in the tropics and subtropics, *E. histolytica* is capable of utilizing host resources and interacting with the microbiota to meet its nutritional and protective needs ^5^. However, though 90% of *E. histolytica* infections are asymptomatic, at least 500 million people are infected worldwide and tens of millions develop amebiasis symptoms, mostly in countries with lower socioeconomic conditions ^3, 6^. Invasive amebiasis causes approximately 100,000 deaths annually, ranking as the second leading cause of death among protozoan parasitosis ^6, 7^.

Amebiasis usually begins with the disruption of intestinal homeostasis, followed by severe tissue destruction and inflammation, eventually leading to massive intestinal ulceration ^8^. In some cases, the parasite enters the bloodstream and causes more fatal extraintestinal abscesses ^9, 10^. Although it is unclear how *E. histolytica* transitions from symbiont to pathogen, human biopsy specimens and experimental animal infections have shown that the early-to-intermediate stages of host intestinal amebiasis are characterized by an intense inflammatory response ^11, 12^. Host innate immune defenses, represented by neutrophils and macrophages, play more important roles at these stages ^13^. Most often, the invading amoebic trophozoites are cleared after an acute inflammatory response, but in a small percentage of individuals, the parasite escapes elimination and intestinal lesions continue to develop ^12^. At this stage, inflammation in the gut is instead attenuated, which may be related to the development of a non-protective adaptive immune response ^12, 14^.

As an indispensable prerequisite for parasite colonization and invasion, *E. histolytica* adherence is mediated by a group of galactose (Gal)- and N-acetyl-D-galactosamine (GalNAc)-inhibitable lectins ^4, 15^. Upon attachment, amoebas induce cell death and tissue invasion through a series of cytotoxic mechanisms, including apoptosis, phagocytosis, and trogocytosis ^9, 16^. In this process, amoebapores, cysteine proteases, amoebic acid vesicle components, and intracellular calcium signaling in trophozoites all play a role ^17–20^. Interestingly, in the initial stage of intestinal amebiasis, though the intestinal epithelial barrier remains intact and no parasite contact with the epithelial cells is observed, moderate inflammation and infiltration of innate immune cells are already present in the subepithelial lamina propria ^14^. This suggests that before *E. histolytica* trophozoites adhere to and kill host cells, some virulence molecules released by the parasite can transit through the epithelium and reach immune cells, acting as pathogen-associated molecular patterns (PAMPs) to initiate intestinal inflammation ^12^. The transepithelial movement of these virulence molecules is contributed by amoebic synthesis and secretion of prostaglandin E2, which disrupts the tight junctions of intestinal epithelial cells by binding to E-prostanoid-4 receptors ^12, 21^. After tight junction disruption, the transfer of *E. histolytica* PAMPs to the basolateral surface of intestinal epithelium can be detected ^22^. Moreover, if a virulence molecule is present in amoebic extracellular vesicles (EVs), it can be endocytosed by intestinal epithelial cells, transported to the basolateral side of epithelial cells through multivesicular bodies, then released into the lamina propria ^23, 24^.

*E. histolytica* uses humans as exclusive natural hosts ^13^. Thus, owing to the lack of suitable animal models to simulate the entire cycle of human disease, the study of amebiasis remains complicated and challenging ^25^. At present, mice are still the most commonly used animals for laboratory studies of *E. histolytica* infection, among which some inbred mouse strains are susceptible to intestinal amebiasis, such as C3H/HeN, C3H/HeJ, and CBA/J ^13, 26, 27^. In recent years, single-cell RNA sequencing (RNA-seq) has become a powerful tool, revolutionizing our understanding of complex biological systems in various infectious diseases at the individual cell level ^28^. This technology allows not only screening for cell types that play an important role in the infection process, but also identifying their interaction patterns with other cells ^29, 30^. However, to date, no host single-cell atlas of clinical samples or animal models has been developed in the field of amebiasis research. In this study, to explore the gene expression characteristics of different host cell types in the acute phase of intestinal amebiasis, *E. histolytica* trophozoites were inoculated into the cecum of C3H/HeNCrl mice and fresh cecal tissues were used for single-cell RNA-seq. After determining the important cell types, underlying mechanisms of their gene expression changes and cell-cell interactions were further explored.

## MATERIALS AND METHODS

### Amoebic strains and cell cultures

Trophozoites of *E. histolytica* isolates, HM-1:IMSS and SAW755CR, were grown under axenic conditions at 36.5°C, in YIMDHA-S medium containing 10% and 15% (v/v) heat-inactivated adult bovine serum (Sigma-Aldrich, St. Louis, MO, USA), respectively.

Human pro-monocytic myeloid leukemia cell lines, U937 and THP-1, were grown in RPMI-1640 medium (#10-040-CV; Corning, Manassas, VA, USA) supplemented with 10% (v/v) fetal bovine serum (HyClone Laboratories, Logan, UT, USA). Human intestinal epithelial cells, Caco-2, were grown in Eagle’s minimum essential medium (MEM; #10-010-CV; Corning, Manassas, VA, USA) containing 20% (v/v) fetal bovine serum, 1% (v/v) MEM non-essential amino acids (Gibco, Grand Island, NY, USA), 1% (v/v) 2 mM L-glutamine (Gibco, Grand Island, NY, USA), and 1% (v/v) 1 mM sodium pyruvate (Gibco, Grand Island, NY, USA). Human HT29-MTX-E12 and mouse RAW264.7 cells were grown in DMEM medium (#10-013-CV; Corning, Manassas, VA, USA) supplemented with 10% (v/v) fetal bovine serum, 1% (v/v) MEM non-essential amino acids, 1% (v/v) 2 mM L-glutamine, and 1% (v/v) 1 mM sodium pyruvate. The above cells were grown at 37°C in a 5% CO_2_ incubator. Human embryonic kidney (HEK) 293 cells were cultured in OPM-293 CD05 medium (81075-001; Opmbiosciences, Shanghai, China) under conditions of 37 °C, 5% CO_2_, and 120 rpm shaking.

After seeding, human U937 and THP-1 monocytes were pretreated with 100 ng/mL phorbol 12-myristate 13-acetate (PMA; Sigma-Aldrich) for 24 h to differentiate them into macrophage-like phenotypes. For the construction of the intestinal epithelial model, human Caco-2 and HT29-MTX-E12 cells were seeded on the apical side of Falcon Transparent Inserts (0.4 μm pore size; 353095; Corning, Tewksbury, MA, USA) in ratios of 3:1 or 9:1. DMEM medium was used across all coculture experiments.

### Animal model for intestinal amebiasis

Seven-week-old male C3H/HeNCrl mice were purchased from the Beijing Vital River Laboratory Animal Technology Company, and maintained under specific pathogen-free conditions. For a more stable induction of intestinal amebiasis, the more virulent SAW755CR strain was used instead of the general HM-1:IMSS strain during intracecal inoculation of *E. histolytica* trophozoites, whose toxicity was maintained by regular passage through the hamster liver (Songlian Experimental Animal Factory, Shanghai, China) as previously mentioned ^31^. Mice in experimental groups were inoculated directly with 100 µL of 1L×L10^6^ axenic trophozoites into the cecum (*n*L≥L3), while mice in control groups received the same volume of YIMDHA-S medium without trophozoites (*n*L≥L3).

Amebiasis severity was evaluated by monitoring animal body weight changes, routine blood tests, and immunohistochemical staining of cecal tissues. At 4, 6, and 8 d post-inoculation, the mice were euthanized, blood samples were collected into 1 mL EDTA K2 tubes, and cecal tissues were harvested for 4% paraformaldehyde fixation. According to the manufacturer’s instructions, routine blood tests were performed using a ProCyte Dx Hematology Analyzer (IDEXX Laboratories, ME, USA), and the proportion of various leukocytes was calculated by ProCyte Dx software version 00-35. Meanwhile, after fixation in paraformaldehyde and paraffin embedding, cecal tissue sections were subjected to hematoxylin and eosin (H&E) staining and immunohistochemical staining. Expression levels of tumor necrosis factor alpha (TNF-α), interleukin (IL)-1β, IL-6, E-cadherin, Claudin-3, and ZO-1 in the cecum were detected.

In strict accordance with the guidelines of the Regulations for the Administration of Affairs Concerning Experimental Animals (1988.11.1), all animal experiments were conducted with the approval of the Institutional Animal Care and Use Committee (permit no. 20160225-097). All efforts were made to minimize animal suffering.

### Other experiments and analyses

All other materials and methods are detailed in Supplemental Methods.

## RESULTS

### Intense inflammatory response in the acute phase of intestinal amebiasis

First, to determine the clinical characteristics of the acute phase of intestinal amebiasis, a mouse model was established by inoculating *E. histolytica* trophozoites into the cecum of seven-week-old male C3H/HeNCrl mice. To ensure the stability of infection, the more virulent *E. histolytica* isolate SAW755CR was used in our animal experiments, as previously mentioned ^31^. Murine body weight, leukocyte proportion, and inflammatory infiltration of the cecal tissue were regularly detected (Figure 1a). After surgery, the infected mice lost weight compared to control-treated mice (Figure 1b). At 6 and 8 d post-inoculation, murine routine blood tests showed not only increased proportions of neutrophils, monocytes, and eosinophils, but also a decreased proportion of lymphocytes, suggesting the presence of inflammation caused by bacterial or parasitic infection (Figure 1c). As the three inflammatory cytokines that have attracted much attention in studies of amebiasis, infiltration of TNF-α, IL-1β, and IL-6 were also detected in the cecal tissue of amoeba-infected mice (Figure 1d) ^8, 13, 32^.

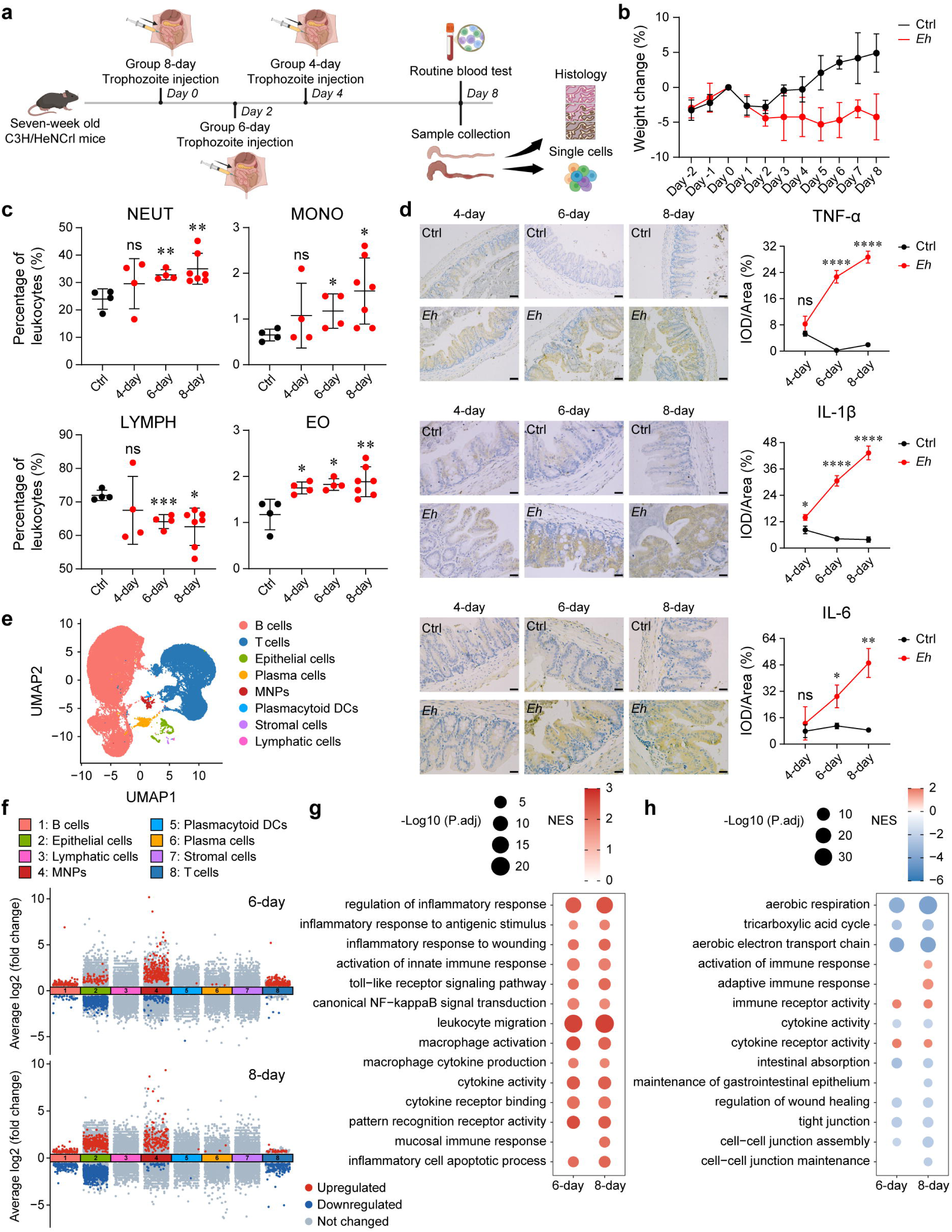

Owing to the obvious pathological changes in mice in Infection 6-day and Infection 8-day groups, single-cell RNA-seq was further performed on their cecal tissues. The experiment consisted of three groups with three mice in each group. Initially, data quality control was conducted to filter out low-quality information after sequencing (Figure S1a). Upon combining all the data, single cells from infected and control-treated samples were clustered and visualized in Figure S1b, and the proportion of cell clusters in each group was shown in Figure S1c. Using previously described cell-specific marker genes (Figure S1d), 68,058 single cells from the ceca of nine different mice were annotated into 15 cell types (Figure 1e) ^29^. Since the more abundant lymph nodes in the cecum and the dominant position of B and T cells in these lymph nodes, the single-cell atlas in our study exhibited higher proportions of the two cell types (Figure S1e) ^33^. Setting |log2FC| value > 0.585 and adjusted p value <L0.05 as the criteria, differentially expressed genes (DEGs) in each cell type in Infection 6-day and Infection 8-day groups were identified (Figure 1f). Detailed gene expression changes of each cell type are shown in Table S1.

Among all cell types, mononuclear phagocytes (MNPs) and epithelial cells showed the greatest differences in terms of DEG numbers and fold changes (Figure 1f). Thus, Gene Set Enrichment Analysis (GSEA) of various Gene Ontology (GO) terms was performed for these two cell types. In the cecum of amoeba-infected mice, MNPs exhibited significant positive enrichment of pro-inflammatory GO terms, which were highly correlated with innate immunity and cytokines (Figure 1g). Epithelial cells showed not only significant negative enrichment of GO terms associated with aerobic respiration and tight junctions, but also significant positive enrichment of immune-related GO terms, suggesting inflammatory injuries of the intestinal epithelium under *E. histolytica* infection (Figure 1h).

### Important role of mononuclear phagocytes in intestinal amebiasis

To further identify the cell types that play an important role in intestinal amebiasis, GSEA of Kyoto Encyclopedia of Genes and Genomes (KEGG) pathways was performed on individual cells and the Amebiasis pathway (mmu05146) was selected for comparison. For the enrichment level of the Amebiasis pathway, every single cell in each group was scored separately and plotted in a Uniform Manifold Approximation and Projection (UMAP) visualization (Figure 2a). As shown by the dashed circles, after *E. histolytica* infection, the most significant enrichment occurred in cecal MNPs. The AUCell scores of the Amebiasis pathway gene set for all cell types were exhibited in Figure 2b. In Infection 6-day and Infection 8-day groups, the scaled scores of MNPs improved the most. Correspondingly, the MNP proportion increased the most after parasitic inoculation (Figure 2c). These results suggested a critical function of host MNPs in intestinal amebiasis.

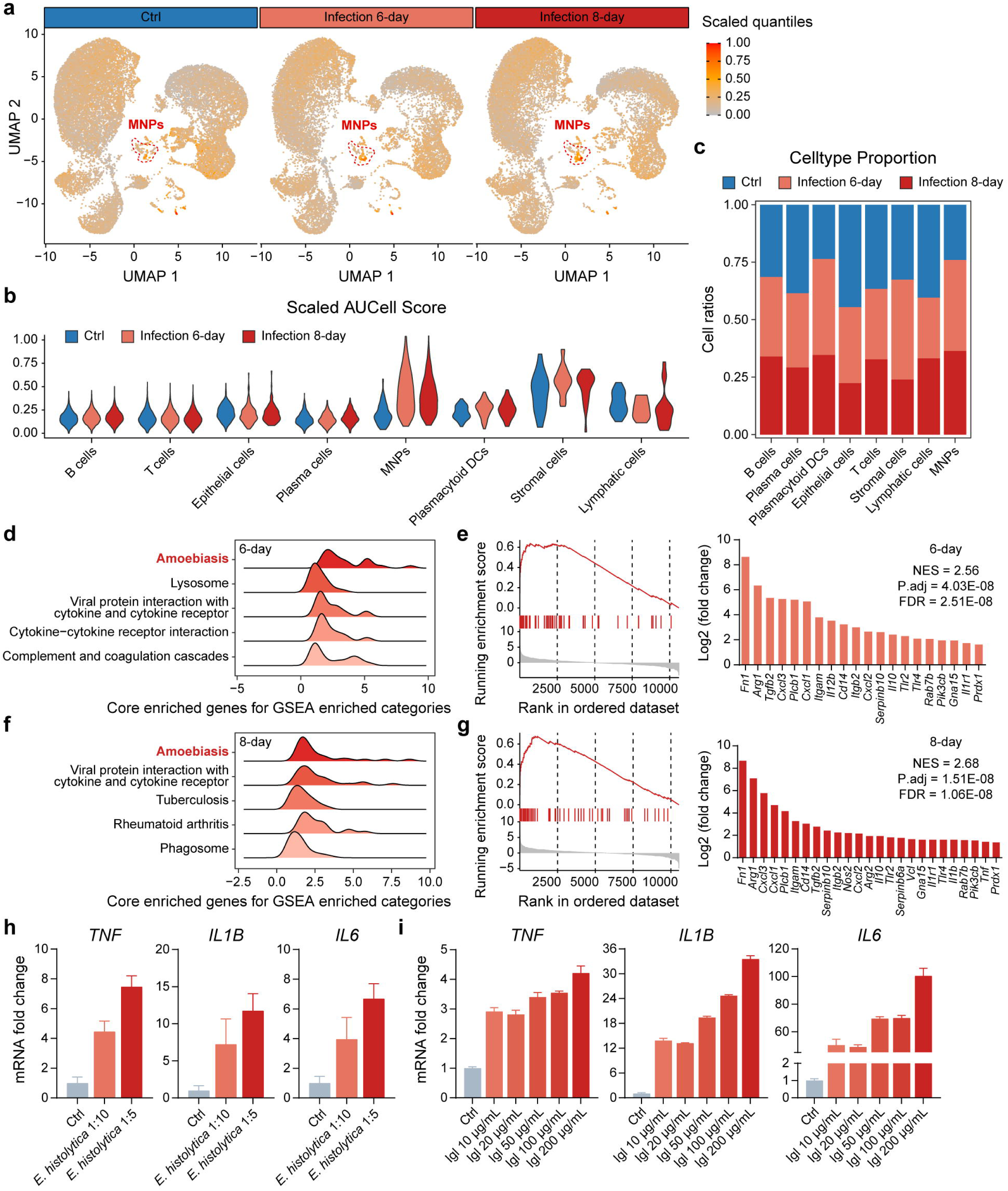

Comparing Infection 6-day group and Ctrl group MNPs, the top five significantly upregulated KEGG pathways in GSEA was shown (Figure 2d). As the most significantly positively enriched one, GSEA of the KEGG Amebiasis pathway was visualized, and all of its core-enriched genes were displayed (Figure 2e). Similarly, comparing Infection 8-day group and Ctrl group MNPs, the top five significantly upregulated KEGG pathways in GSEA was shown in Figure 2f, while GSEA plot of the most significantly positively enriched Amebiasis pathway and its core-enriched genes were displayed in Figure 2g. Under GSEA, all significantly altered KEGG pathways of cecal MNPs are detailed in Figure S2a to S2d. The filter criteria were set as |NES| > 1, adjusted p valueL<L0.05, and FDR <L0.25.

In KEGG enrichment analysis of MNPs, the Amebiasis pathway was the most upregulated both 6 and 8 d post-inoculation, suggesting that intestinal inflammation in mice was more likely to be induced directly by *E. histolytica* trophozoites than indirectly by the imbalanced gut microbiota caused by parasite colonization. From the perspective of human cell line, the pro-inflammatory effect of *E. histolytica* trophozoites on PMA-differentiated U937 monocytes was further demonstrated (Figure 2h). As the intermediate subunit of *E. histolytica* Gal/GalNAc lectin, Igl is an important molecule on the amoebic cell surface that contributes to parasite adherence ^34^. Native Igl protein could also act as a PAMP to stimulate human U937 cells for inflammatory signal initiation (Figure 2i).

### *E. histolytica* Igl induces an inflammatory response in macrophages through its C3 segment

The C3 region of *E. histolytica* Igl not only has the most conserved propensity among various Igls of *Entamoeba* species, but also plays a potential role in the carbohydrate recognition of parasite adherence, making it more likely to participate in host immune recognition as an important PAMP segment ^8^. Using lipopolysaccharide (LPS) and IL-4 as controls, eukaryote-expressed *E. histolytica* Igl-C3 could activate host macrophages and differentiate them into a pro-inflammatory phenotype, but changes in their cytokine transcription levels were smaller than those of LPS stimulation (Figure 3a). In addition to TNF-α, IL-1β, and IL-6, expressions of IL-10, IL-23, and TGF-β1, which are often mentioned in studies of amebiasis, were also examined ^32, 35^. To detect cytokine production and release, western blotting of U937 cells and enzyme-linked immunosorbent assay (ELISA) of cell culture supernatants were performed respectively (Figure 3b and 3c). After Igl-C3 stimulation, cytokine expression of PMA-differentiated human THP-1 cells and mouse RAW264.7 cells was further examined to confirm this pro-inflammatory effect (Figure S3a and S3b).

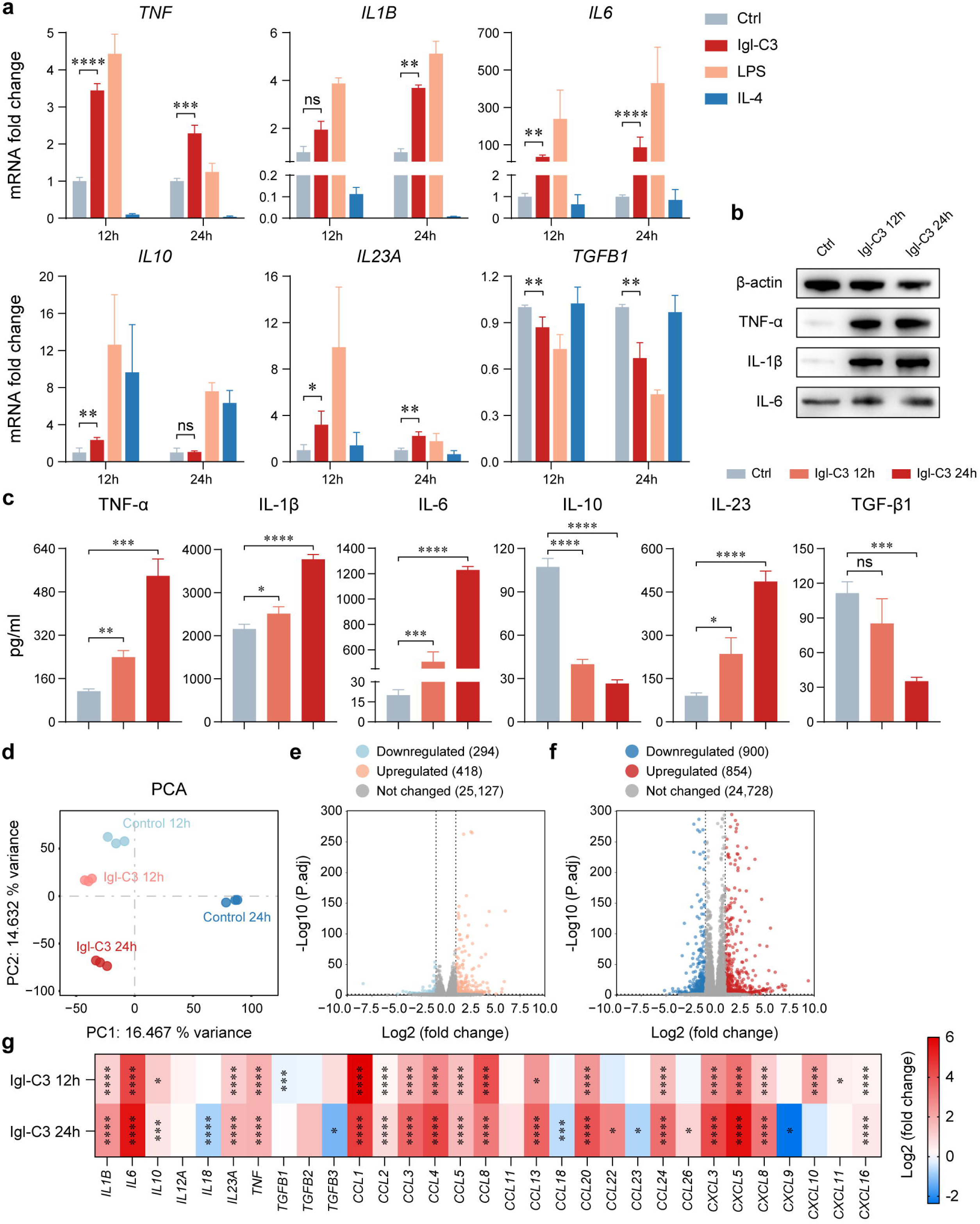

To better understand the pro-inflammatory effect of Igl-C3, transcriptomic sequencing of differentiated U937 cells was performed after eukaryote-expressed Igl-C3 stimulation for 12 or 24 h. Gene expression levels of each sample were shown in Figure S4a, and correlations between the gene expression profiles of any two samples from four groups were shown in Figure S4b. In the principal component analysis, stratification between the Igl-C3-treated 24 h group and the Control 24 h group was more obvious than that between the Igl-C3-treated 12 h group and the Control 12 h group (Figure 3d). With increased time of Igl-C3 stimulation, the DEG number in macrophages increased and DEG fold changes were more significant (Figure 3e and 3f). At different stimulation time points, changes in cytokine expression detected by the transcriptome were detailed in Figure 3g, verifying the pro-inflammatory function of Igl-C3.

In GO enrichment analysis of the transcriptome, human U937 cells exhibited stronger immune responses and cytokine activity after *E. histolytica* Igl-C3 stimulation (Figure S4c and S4d). Comparing the Igl-C3-treated 12 h group and Control 12 h group macrophages, the top 20 significantly altered KEGG pathways and their rich factors were shown in Figure S5a. Interestingly, significance of the change in the KEGG Amebiasis pathway (hsa05146) ranked first among terms of infectious diseases, suggesting that the C3 region of *E. histolytica* Igl might be an important characteristic PAMP in amebiasis. The DEG changes in the Amebiasis pathway were further detailed in Figure S5b. Similarly, comparing the Igl-C3-treated 24 h group and Control 24 h group macrophages, significance of the change in the KEGG Amebiasis pathway also ranked first among terms of infectious diseases (Figure S5c). After stimulating macrophages for 24 h, the DEG changes in the Amebiasis pathway were detailed in Figure S5d.

### Igl-C3 indirectly impairs intestinal epithelium integrity by inducing host inflammation

Single-cell RNA-seq of mouse cecal tissues revealed that besides MNPs, epithelial cells were another cell type with significant gene expression changes (Figure 1f). The upregulation of DEGs was most significant in MNPs, while the downregulation of DEGs was most significant in epithelial cells. To explore whether DEG changes in these two cell types were correlated, the interactions between different cell types in the cecum were analyzed using the CellChat database. In the Ctrl group, no cellular communication from MNPs to epithelial cells was detected, but such interactions emerged 6 and 8 d after parasitic inoculation (Figure 4a). With the persistence of *E. histolytica* infection, the interaction number from MNPs to epithelial cells continued to increase (Figure 4b). At different time points after amoebic infection, detailed changes in the cell-cell interactions between MNPs and epithelial cells were revealed in Figure 4c, being associated with immune regulation, host defense, and transduction of inflammatory signals.

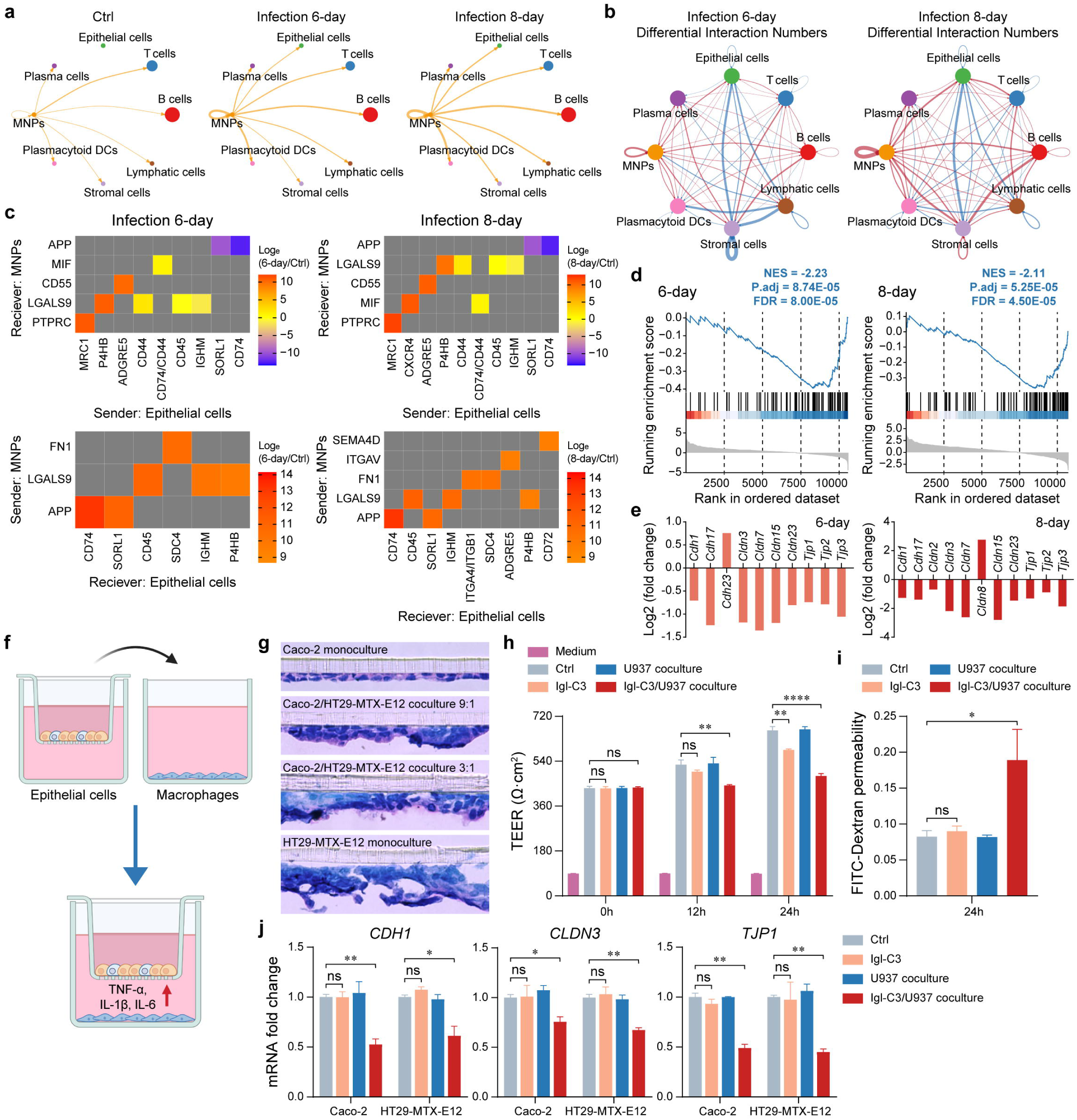

In addition to intense inflammation, another characteristic of intestinal amebiasis is severe destruction of the gut tissue ^8^. In our study, the GSEA plots exhibited significant negative enrichment of GO Tight Junction term (GO:0070160) in the cecal epithelial cells of amoeba-infected mice (Figure 4d). The *Cdh*, *Cldn*, and *Tjp* gene families play important roles in maintaining the tight junctions of intestinal epithelium, and their gene expressions were also mostly decreased after *E. histolytica* trophozoite infection (Figure 4e) ^3, 36–38^. For each cell type in the single-cell RNA-seq data, incoming signaling patterns of cellular communication in different groups were shown in Figure S6a, and outgoing signaling patterns of cellular communication in different groups were shown in Figure S6b. In terms of the CDH- and CLDN-related pathways, the intensity of signal sending and receiving in epithelial cells was significantly upregulated, suggesting ongoing changes in tight junctions and barrier functions. Based on their importance in studies relevant to intestinal tight junctions, the *Cdh*, *Cldn*, and *Tjp* gene family members, *Cdh1*, *Cldn3*, and *Tjp1*, respectively, were selected for further detection ^3, 36, 39^. The decreased expression of E-cadherin (*Cdh1*), Claudin-3 (*Cldn3*), and ZO-1 (*Tjp1*) in mouse cecal tissues was examined by immunohistochemistry to verify the single-cell RNA-seq results (Figure S7a to S7c).

Subsequently, to determine whether *E. histolytica* Igl-C3 contributes to this injury of intestinal epithelium, a three-dimensional triple culture system was established for mimicking the structure of epithelial cell monolayer and lamina propria macrophages (Figure 4f). The epithelial cell monolayer consists of Caco-2 and HT29-MTX-E12 cells, whose common ratios for simulating the intestinal epithelium are 3:1 and 9:1 ^40, 41^. In the present study, after comparing mucus production, a 3:1 ratio was used for follow-up experiments (Figure 4g). Upon Igl-C3 treatment for 24 h, transepithelial electronic resistance (TEER) of the epithelial cell monolayer decreased slightly, whereas in the presence of macrophages, Igl-C3 incubation resulted in a much more significant TEER reduction (Figure 4h). Similarly, in the presence of macrophages, Igl-C3 treatment significantly increased the permeability of epithelial cell monolayers to FITC-Dextran (Figure 4i). In response to macrophage inflammation induced by Igl-C3, expressions of tight junction-related *CDH1*, *CLDN3*, and *TJP1* genes were all reduced in epithelial Caco-2 and HT29-MTX-E12 cells (Figure 4j).

### Macrophages recognize Igl-C3 via TLR4 and initiate the TLR4/MyD88/NF-**κ**B inflammatory signaling pathway

Our results suggested that Igl-C3 could act as a PAMP to induce an inflammatory response in macrophages and indirectly impairs intestinal epithelium integrity through inflammatory cytokines released by macrophages. Since the pathogenic effect of Igl-C3 in intestinal amebiasis was mainly reflected in its interaction with macrophages, the underlying mechanism of such interaction was subsequently explored. In the transcriptomic sequencing of Igl-C3-treated U937 cells, the interaction network between the top 20 KEGG pathways with the most significant upregulation indicated that changes in the Amebiasis pathway might be related to toll-like receptor (TLR) signals (Figure 5a and 5b). As the two TLRs being reported to identify *E. histolytica* PAMPs, the responses of TLR2 and TLR4 to Igl-C3 were detected by preincubation with the corresponding inhibitors (Figure 5c) ^42, 43^. Macrophages were demonstrated to recognize C3 segment of *E. histolytica* Igl via TLR4.

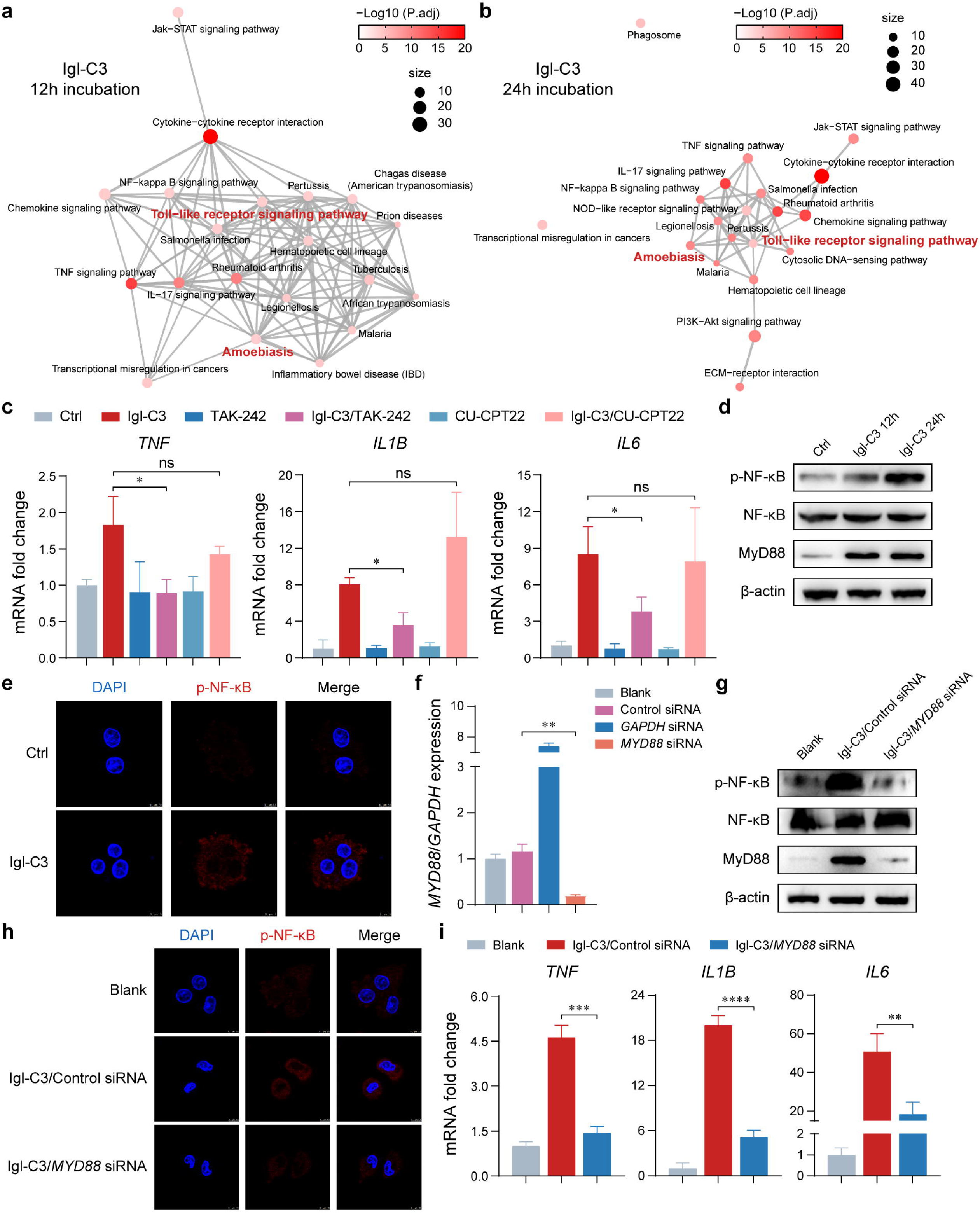

After the recognition of Igl-C3 by TLR4, inflammatory signals within host macrophages were found to be initiated by the classical TLR4/MyD88/NF-κB signaling pathway through western blotting and laser confocal microscopy (Figure 5d and 5e) ^44^. Upon 24 h of Igl-C3 treatment, the expression of MyD88 in macrophages increased, and the expression of phospho-NF-κB was upregulated relative to total NF-κB, exhibiting a tendency to enter the nucleus. Small interfering RNA (siRNA) treatment was further performed to silence the *MYD88* gene in macrophages for verification (Figure 5f). After siRNA treatment, the TLR4/MyD88/NF-κB signaling pathway was inhibited accordingly (Figure 5g and 5h), and expressions of pro-inflammatory cytokines were also significantly reduced (Figure 5i).

In the single-cell RNA-seq data, comparing the Infection 6-day group and Ctrl group MNPs, a GSEA plot of the KEGG Toll-like receptor signaling pathway (mmu04620) and its core-enriched genes were displayed in Figure S8a. Similarly, comparing the Infection 8-day group and Ctrl group MNPs, that GSEA plot and core-enriched genes were displayed in Figure S8b. At different time points upon amoebic infection, the gene sets of Toll-like receptor signaling pathway all showed a significant positive enrichment, and expressions of *Tlr4*, *Myd88*, and *Nfkbia* genes increased, suggesting that the TLR4/MyD88/NF-κB signaling pathway might be active and crucial to the pro-inflammatory process of macrophages during intestinal amebiasis.

### Igl-C3 binds to the TLR4 co-receptor MD2 on the surface of macrophages to activate inflammatory signals

To determine how TLR4 recognized Igl-C3 on the surface of host macrophages, inhibitor and siRNA treatments were first performed on human U937 cells targeting the TLR4 co-receptor MD2. As an indispensable accessory protein of TLR4, MD2 can bind to it and recruit the adaptor protein MyD88, producing a large number of pro-inflammatory cytokines ^45^. After preincubation with the inhibitor, the TLR4/MyD88/NF-κB inflammatory signaling pathway of the macrophages was inhibited, and the transcription and translation levels of pro-inflammatory cytokines were decreased (Figure 6a and 6b). The release of pro-inflammatory cytokines into the cell culture supernatant was also significantly reduced (Figure 6c). Targeting MD2 (*LY96*), the siRNA treatment on human U937 cells was shown in Figure 6d and 6e. After silencing the *LY96* gene, expressions of pro-inflammatory cytokines in macrophages decreased significantly (Figure 6f).

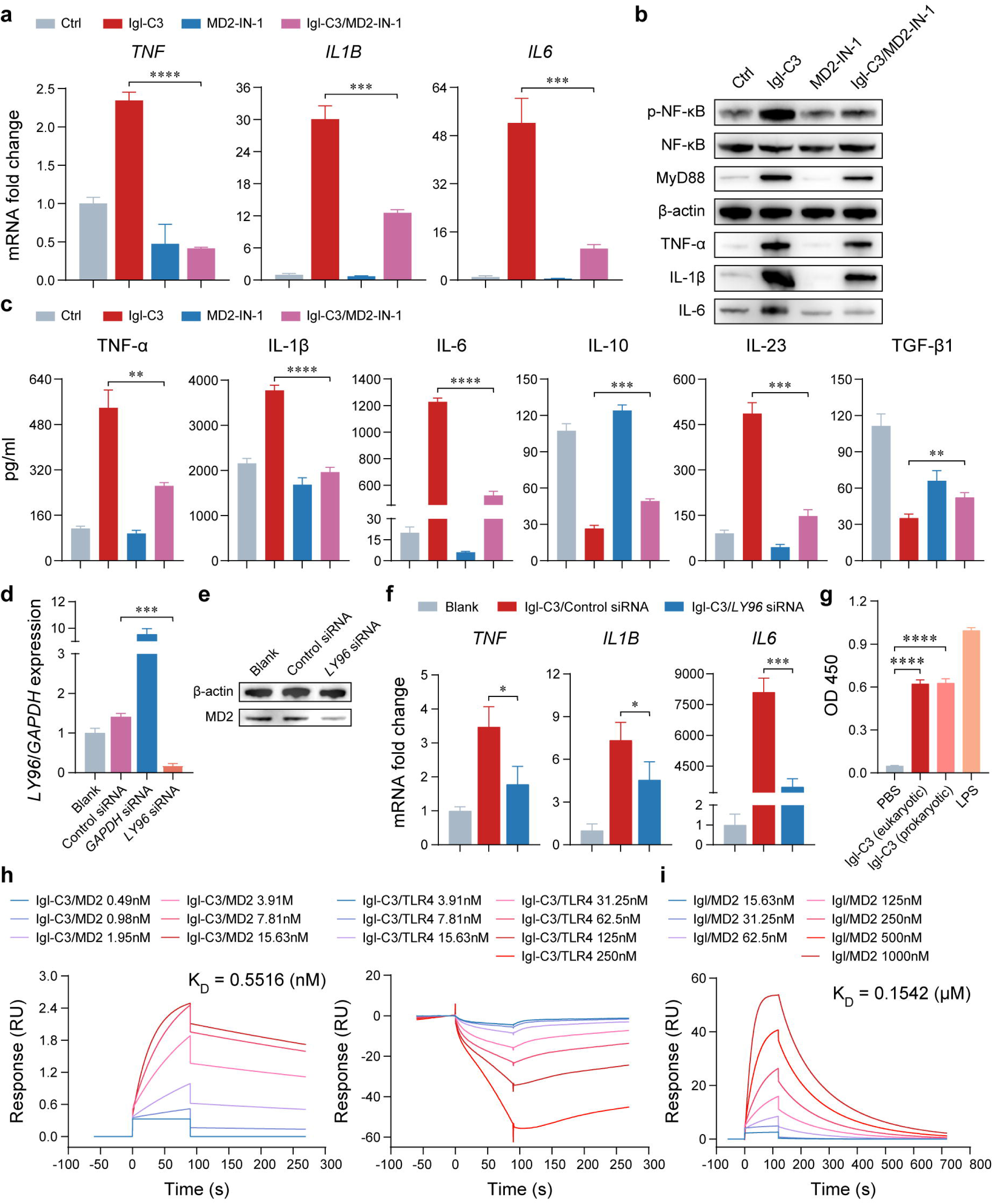

In the ELISA experiments, both eukaryotic Igl-C3 and prokaryotic Igl-C3 exhibited potential binding to human MD2, but the strength was lower than that of LPS (Figure 6g). Subsequently, Biacore analysis was performed to detect the protein affinities of eukaryotic Igl-C3/human MD2 and eukaryotic Igl-C3/human TLR4. As shown in Figure 6h, Igl-C3 could bind to MD2, but could not directly bind to TLR4. Protein affinity between native *E. histolytica* Igl and human MD2 was also demonstrated, indicating the presence of such stimulation in the clinical infection state (Figure 6i).

### *E. histolytica* Igl diffuses along with amoebic EVs to stimulate host macrophages

At the end of the present study, EV-related experiments were conducted to explore how *E. histolytica* Igl contacts host macrophages to induce pro-inflammatory signals in intestinal amebiasis. The presence of Igl in the amoebic EVs was first confirmed by western blotting (Figure 7a). Subsequently, *E. histolytica* trophozoites were cocultured with host macrophages, and quantitative proteomics was applied to compare changes in protein composition and function between the macrophage coculture group and control culture group amoebic EVs. As shown in Figure 7b, a total of 1,693 different proteins were annotated, and 1,072 of which were shared between the two groups. The top 20 most abundant proteins in the control culture group amoebic EVs were exhibited in Figure S9a, and the top 20 most abundant proteins in the macrophage coculture group amoebic EVs were exhibited in Figure S9b. Within each group, the fractions of total protein abundance across independent biological replicates were generally consistent (Figure 7c). Meanwhile, high correlations existed among samples in each group (Figure 7d).

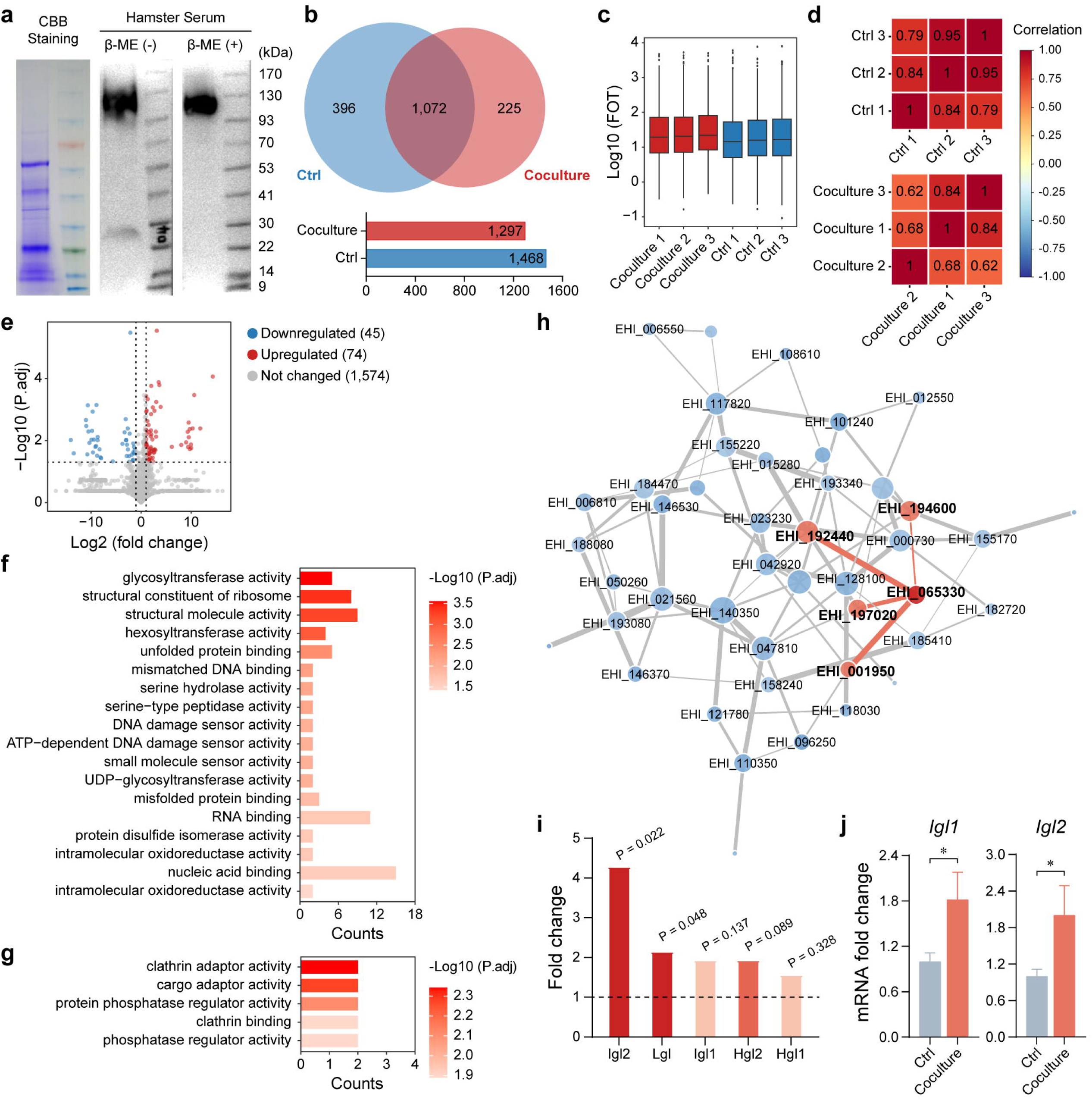

After coculture with host macrophages, 74 proteins were upregulated and 45 proteins were downregulated in amoebic EVs (Figure 7e). To identify the variations in protein function, changes of terms under Molecular Function in the GO enrichment analysis were compared between the macrophage coculture group and control culture group amoebic EVs (Figure 7f and 7g). Upon coculture, the molecular function of proteins in amoebic EVs showed a more obvious upregulation trend, mostly related to binding and catalysis. Comparing macrophage coculture group and control culture group amoebic EVs, the interaction network of upregulated differential proteins was exhibited in Figure 7h, and that of downregulated differential proteins was exhibited in Figure S9c. In the two subtypes of *E. histolytica* Igl, the content of Igl2 in amoebic EVs was significantly increased, which was potentially associated with heat shock protein (HSP) 70, molybdenum cofactor sulfurase, and signal peptidase complex catalytic subunit ^34^. Except for Igl2, the content of Lgl in amoebic EVs also increased significantly after coculture with host macrophages (Figure 7i). Finally, upon incubation with macrophages, expressions of both Igl1 and Igl2 in *E. histolytica* trophozoites were upregulated, consistent with the content change trend of these proteins in amoebic EVs (Figure 7j).

## DISCUSSION

The gut mucosal immune system consists of epithelial cells, lamina propria, and lymph nodes, constituting a protective barrier for intestinal tract integrity and having indispensable contributions in resisting pathogen infections ^46^. Host innate immune defenses in this system, represented by neutrophils and macrophages, always play an important role during *E. histolytica* infection ^13^. Recruited by inflammatory cytokines released by epithelial cells and several immune cells, neutrophils migrate from the circulation to infection sites, where they show multiple antiamoebic functions such as degranulation, phagocytosis, and formation of neutrophil extracellular traps (NETs) ^12, 47, 48^. However, the role of neutrophils is double-edged, as they are also induced to die by *E. histolytica* trophozoites, in turn promoting inflammation and tissue damage through the release of their lytic products ^49, 50^. Similarly, macrophages perform complicated and two-sided functions during amebiasis. Host macrophages recognize several PAMPs of the amoeba via TLR2 and TLR4, then produce various inflammatory cytokines such as TNF-α, IL-1β, and IL-6 ^42, 43, 51^. Nitric oxide (NO) is an anti-amoebic agent that inhibits essential enzymes involved in *E. histolytica* metabolism ^52^. On the one hand, activated macrophages can mediate the amoebicidal effect of interferon-gamma, producing large amounts of NO from L-arginine through NO synthase ^13^. On the other hand, the abundant inflammatory cytokines released by macrophages not only conduce to the recruitment of innate immune effectors and the initiation of adaptive immune responses, but also aggravate the damage of intestinal tissue ^53^. During the formation of amoebic liver abscess, monocytes have already been demonstrated to cause severe liver damage by releasing microbicidal factors ^54^. In this study, based on single-cell RNA-seq results, we focused on the significant pro-inflammatory effects of *E. histolytica* and its virulence molecule, Igl, on macrophages. Notably, host adaptive immunity appears to lack protection against *E. histolytica* infection, as cellular immune responses are suppressed during the acute phase of intestinal amebiasis, whereas humoral immunity has little influence on disease progression, outcome, or recurrence ^55, 56^.

In the present study, acting as a PAMP, the pro-inflammatory effects of *E. histolytica* Igl on host macrophages through its C3 region were systematically investigated. On the amoebic cell surface, Igl plays an important role as an intermediate subunit of Gal/GalNAc lectin during parasite adherence ^34^. PAMPs are mostly conserved molecules on the surface of pathogens, among which adhesion-related proteins have been extensively studied in multiple pathogens due to their highly conserved propensities and spatial structures easier to contact host cells ^57, 58^. Adhesion proteins FadA (binding E-cadherin and VE-cadherin) and Fap2 (binding Gal/GalNAc) exist on the cell surface of *Fusobacterium nucleatum*, not only helping the bacteria attach to intestinal epithelium, but also inducing the host to produce inflammatory cytokines and recruit immune cells ^59–61^. On the cell surface of *Escherichia coli*, the adhesion protein FimH (binding mannose) can be recognized by TLR4 of host cells, and then activate their TLR4/MyD88/NF-κB signaling pathway to express various inflammatory cytokines ^62^. Recognized by TLR7 of host cells as a PAMP, the hemagglutinin protein (binding sialic acid) of the influenza virus can activate inflammatory signals and trigger an antiviral innate immune response ^63, 64^. Likewise, on the surface of *E. histolytica*, Hgl is the heavy subunit of Gal/GalNAc lectin (binding Gal/GalNAc), which can stimulate intestinal epithelial cells via TLR2 and TLR4, leading to the activation of intracellular NF-κB signaling and the release of pro-inflammatory cytokines ^51, 65^. However, though it is similar in function and distribution to Hgl, the binding of *E. histolytica* Igl to host cell receptors has not been reported ^8^. Except for these Gal/GalNAc lectin component proteins, lipophosphopeptidogylcan is another important molecule on the amoebic cell surface, which can be recognized by TLR2 and TLR4 of macrophages and activate inflammatory signals ^13, 42^. In the amoebic cytoplasm, peroxiredoxin and *E. histolytica* DNA are also effective PAMPs to a certain extent ^43, 66^. Various amoebic PAMPs, including Igl, might contribute greatly to the inflammatory formation of intestinal amebiasis ^12^.

Under the mucous layer, epithelial cells and innate immune cells form a second layer of protection through complex interactions, jointly participating in the immune surveillance of host intestine ^46, 67^. In the gut mucosal immune system, epithelial cells not only directly resist pathogens as a physical barrier, but also send signals to nearby immune cells by producing different cytokines ^46^. Here, oppositely, through single-cell RNA-seq and validation using a three-dimensional triple culture system, *E. histolytica* Igl-C3 was found to indirectly impairs intestinal epithelial integrity by inducing macrophage inflammation. In other inflammatory diseases of the gut, this important role of macrophages and their interactions with intestinal epithelial cells has already been elucidated ^68^. As an example, in inflammatory bowel disease (IBD), activated macrophages typically infiltrate the intestine in large numbers ^67, 69^. In both Crohn’s disease and ulcerative colitis, inflammatory cytokines including TNF-α, IL-1β, and IL-6 released by the macrophages are identified as key factors causing intestinal epithelial damage ^68, 70–72^. For epithelial cells, pro-inflammatory changes in macrophages can promote their expression of claudin-2, a pore-forming tight junction molecule, and trigger the internalization and degradation of barrier-sealing molecules such as E-cadherin and claudin-4, ultimately leading to intestinal barrier leakage ^67, 68^. Interestingly, intestinal amebiasis and IBD not only share similar clinical manifestations, but also have genetic links in susceptibility and sometimes co-occur in a single patient ^8, 73, 74^. In our single-cell RNA-seq and bulk transcriptomic sequencing data, KEGG enrichment analysis of both mouse MNPs in the amoeba-infected animal ceca and human macrophages after *E. histolytica* Igl-C3 stimulation revealed significant positive enrichment of the Inflammatory bowel disease pathway (Figure 5a and S2a). These similar mechanisms of intestinal inflammation formation and epithelial damage might partly explain such curious relationship between the two diseases. Except for IBD, aflatoxin B1 in food can exert non-carcinogenic toxicities, over-recruiting monocytes and macrophages to induce severe colitis in BALB/c mice ^75^. Subsequently, activated macrophages secrete abundant inflammatory cytokines that damage the colonic epithelium. Moreover, during *Clostridioides difficile* infection, the bacteria can induce host macrophages to release inflammatory cytokines through its production of toxins A and B, inhibiting the repair of intestinal epithelial damage and resulting in severe diarrhea ^76^.

Communication between parasites and host immune system involves EVs derived from both pathogens and host cells, which transmit information from one to the other in ways such as proteins and RNA ^77^. During *E. histolytica* infection, more than 200 amoebic proteins were detected in the extracellular environment, being involved in parasite colonization, cyst formation, and evasion of the host immune system ^78^. However, many of these proteins do not contain secretory signaling sequences, indicating their departure from amoebic trophozoites via non-classical secretory pathways such as the secretion of EVs ^79^. *E. histolytica* EVs are approximately 80 to 400 nm in diameter and have morphological features typical of EVs from other systems ^24, 80, 81^. Through omics detection of the RNA and proteins within them, amoebic EVs were found to contributed greatly to parasite communication, development, and pathogenicity 79-81_._

As the two innate immune cell types playing important roles in amebiasis, interactions of *E. histolytica* EVs with host neutrophils and macrophages have already been preliminarily studied ^24, 81, 82^. Both murine bone marrow-derived monocytes and human THP-1 macrophages can internalize and be activated by amoebic EVs, upregulating multiple inflammatory signaling pathways and increasing pro-inflammatory cytokine expressions ^81, 82^. During hepatic amebiasis, parasite EVs might drive the monocyte-mediated immunopathology, which induces a stronger pro-inflammatory effect in male-derived monocytes than in female-derived ones ^81^. Likewise, in a recent study, after coculture of *E. histolytica* trophozoites with host neutrophils, amoebic EVs were collected and their protein composition and biological function were examined ^24^. Probably by carrying and transporting reactive oxygen species, these EVs can exert an immunomodulatory effect on activated neutrophils, downregulating their respiratory burst and NET production ^24^. As a contrast, in our study, the parasite was cocultured with host macrophages, and quantitative proteomic sequencing was then performed on the amoebic EVs. Impressively, whether cocultured with host macrophages or neutrophils, *E. histolytica* EVs exhibited not only upregulated binding and catalysis functions in GO enrichment analysis, but also increased the relative abundance of Gal/GalNAc lectin component proteins (Figure 7f and 7i) ^24^. Besides, the most abundant proteins in these studies were very similar, and some of which were potentially associated with Gal/GalNAc lectins. For example, abundant aldehyde-alcohol dehydrogenase and myosin heavy chain proteins in EVs can, together with Gal/GalNAc lectins, form the uroid region during capping, contributing to the removal of antibodies and other opsonizing molecules from the parasite surface ^24, 83, 84^. Combined with such EV omics results of parasite coculture with macrophages or neutrophils, Gal/GalNAc lectin was suggested to play a more complex and critical role in the host innate immune defense. Within the first five minutes of trophozoite incubation with host cells, the transfer of Gal/GalNAc lectins to target cells has been previously reported, which could also occur through EVs ^85^. Moreover, our study found that the increased Igl2 level in amoebic EVs has a potential functional association with HSP 70. Interestingly, the content of HSP 70 correlate with *E. histolytica* virulence, and the relationship between these two proteins deserves further exploration ^86, 87^.

Overall, using an infection model of intracecal inoculation, the present study identified the gene expression characteristics of different host cell types during the acute phase of intestinal amebiasis. To our knowledge, this is the first report showing a host single-cell atlas in the field of amebiasis research. Further, we proposed that *E. histolytica* Igl diffuses along with amoebic EVs and contacts host macrophages, binding to the TLR4 co-receptor MD2 through its C3 region and initiating the TLR4/MyD88/NF-κB inflammatory signaling pathway in host cells (Figure 8). By inducing macrophage inflammation and cytokine production, *E. histolytica* can indirectly impair the integrity of intestinal epithelium through Igl protein. In response to the parasite invasion during intestinal amebiasis, our results provide new insights elucidating the inflammatory formation and epithelial damage of gut mucosal immune system, and also conduce to the identification of potential drug targets for these amoebas. However, this study still has some limitations. To date, colon is the most common site for *E. histolytica* infection, while the relatively mature intracecal inoculation model is basically used in animal experiments, leading to insufficient simulation of intestinal amebiasis ^12, 13^. To mimic amoebic colitis, a mouse model was used by ligating the animal colon at its proximal end with a length of approximately 2 cm and inoculating *E. histolytica* trophozoites into the colonic loop ^88^. This colonic loop model is currently applied to short-term infections at 3 h, but improvements may extend its duration of infection and better simulate intestinal amebiasis. Moreover, owing to the ability of accumulating polyploid cells in their proliferative phase, gene silencing of wild-type Igl and alternative expression of C3-deficient Igl in *E. histolytica* trophozoites are difficult to achieve ^8^. In this study, no Igl-edited parasites were used to verify the pro-inflammatory effects of Igl-C3 on host macrophages, which is also the direction of our follow-up efforts.

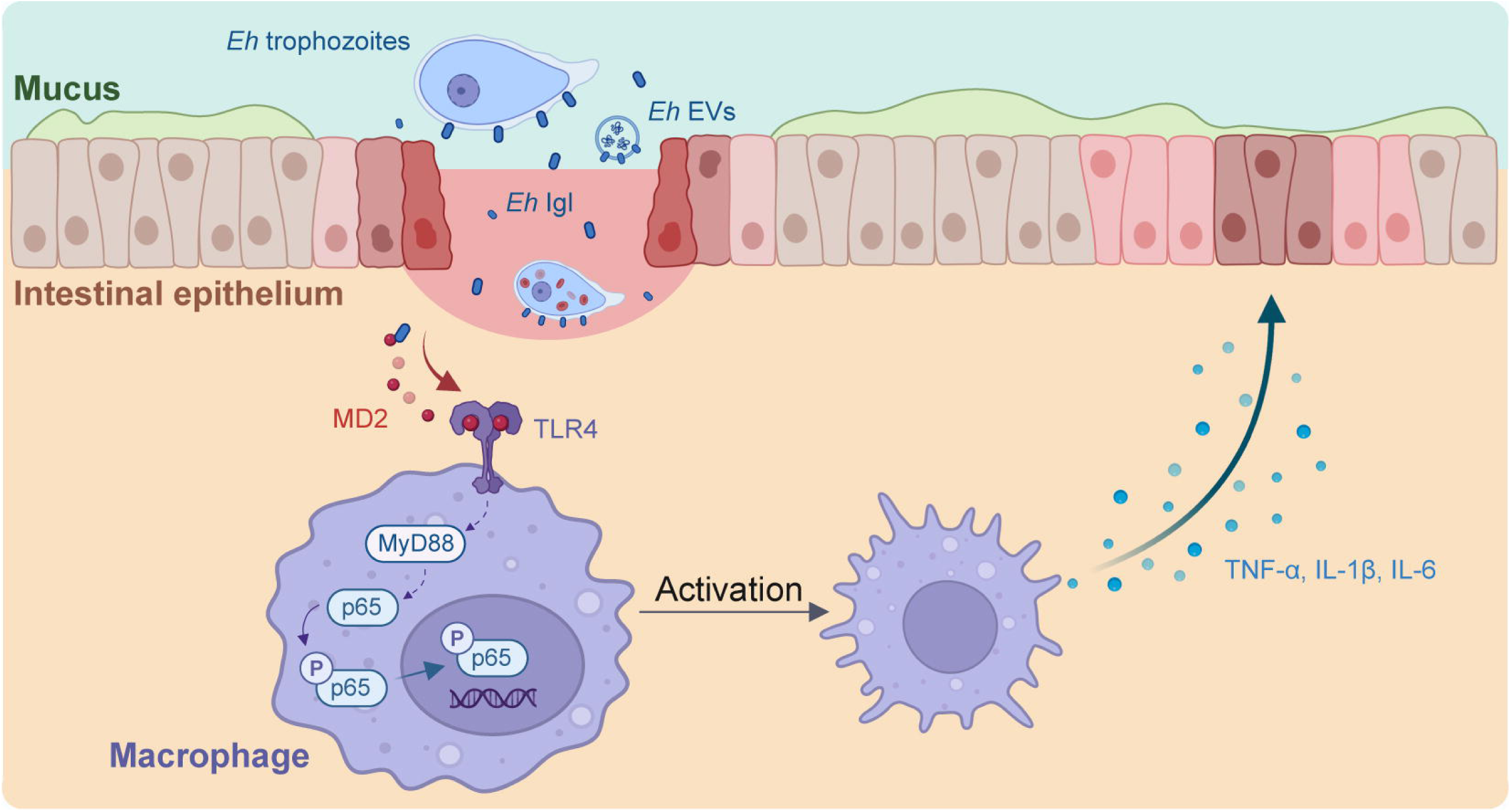

## DATA AVAILABILITY

The datasets generated during this study can be obtained by contacting the corresponding author.

## AUTHOR CONTRIBUTIONS

HZ: Writing - original draft, Methodology, Investigation. SP: Methodology, Investigation. MF: Writing - review & editing, Methodology, Investigation. DD: Investigation. YZ: Investigation. RZ: Investigation. WL: Investigation. XC: Writing - review & editing, Conceptualization. All co-authors revised the manuscript and agreed to its publication.

## ACKNOWLEDGMENTS

We thank all members of the laboratory for fruitful discussions and constructive suggestions.

## ADDITIONAL INFORMATION

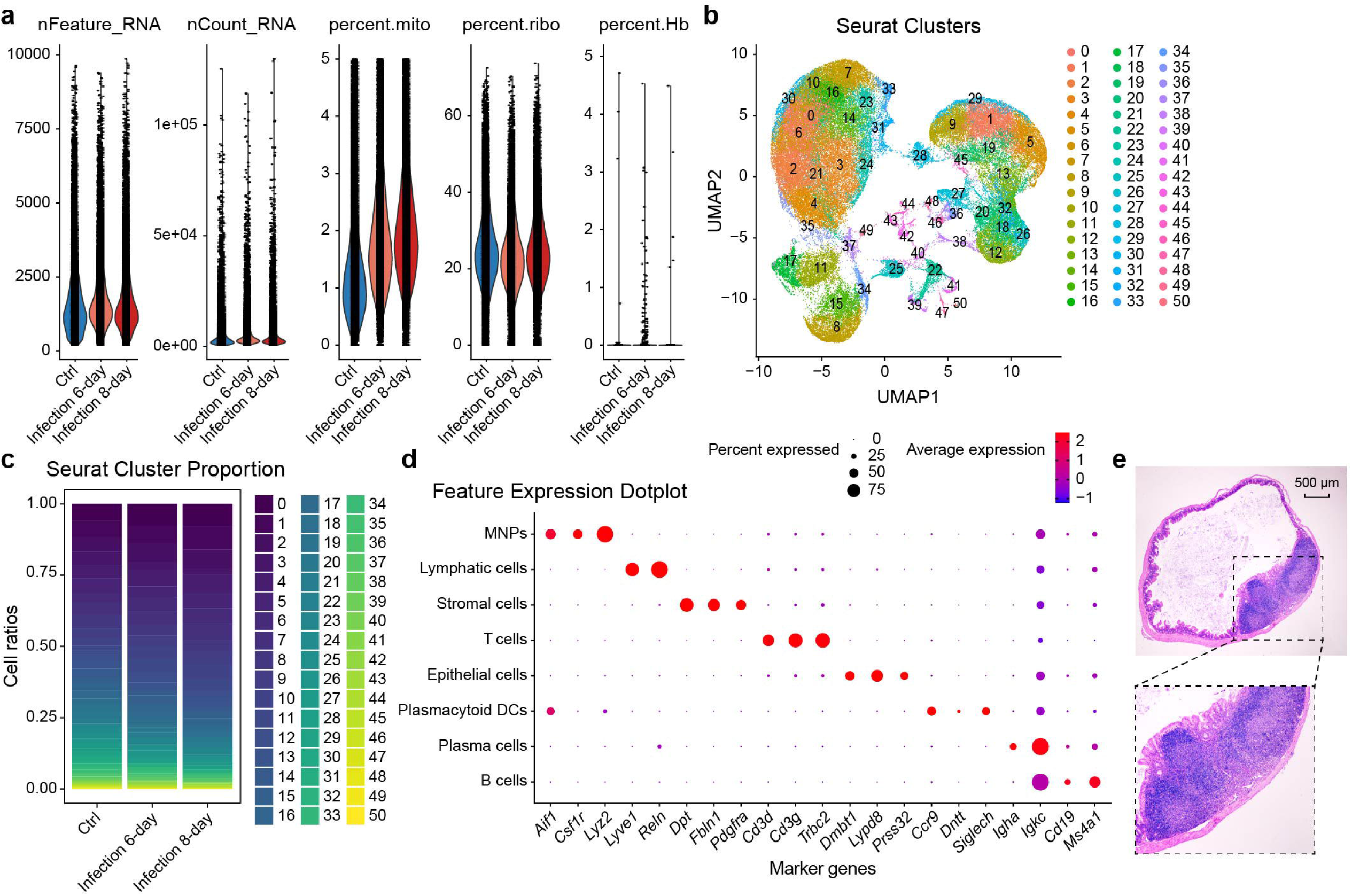

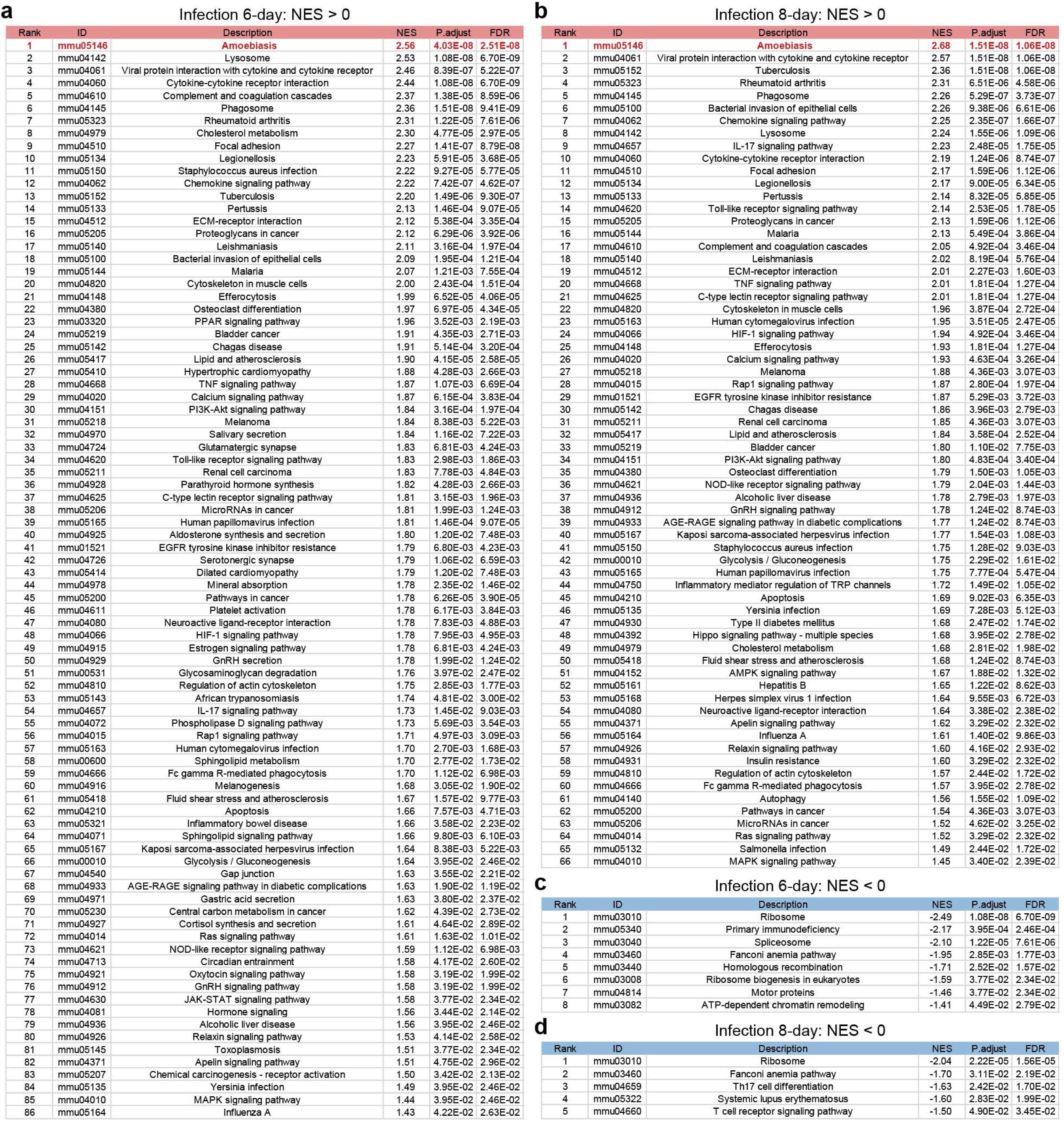

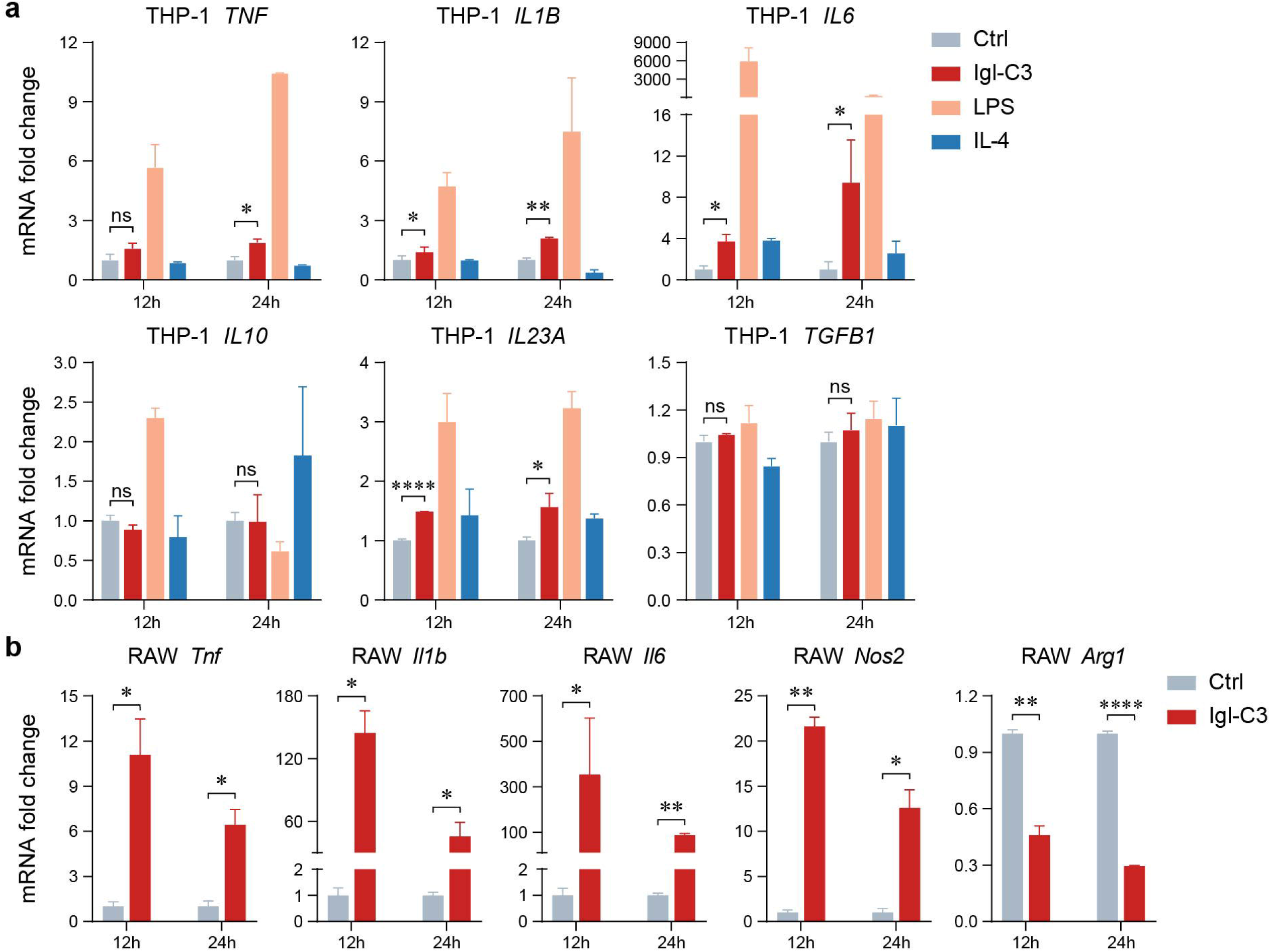

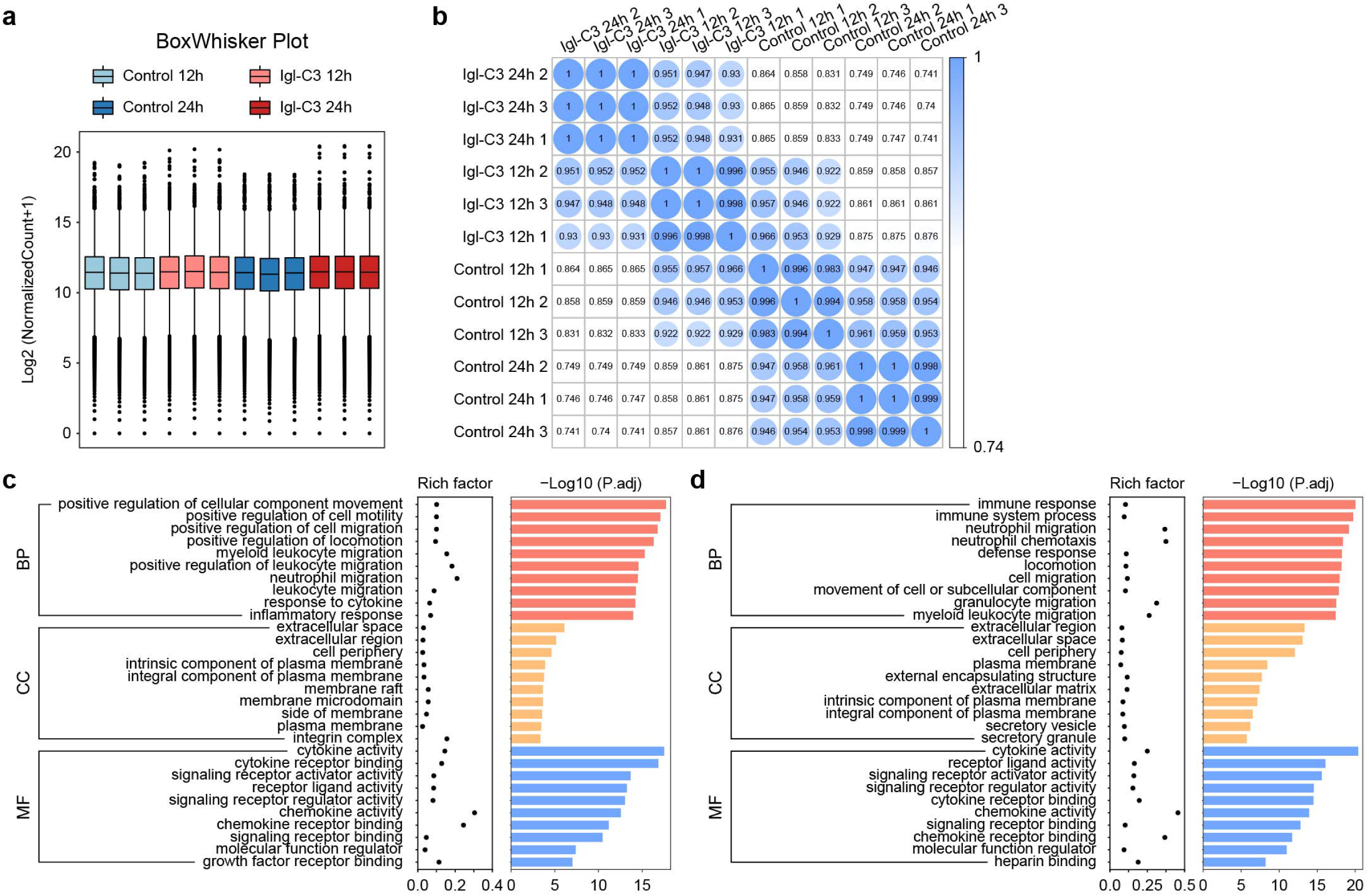

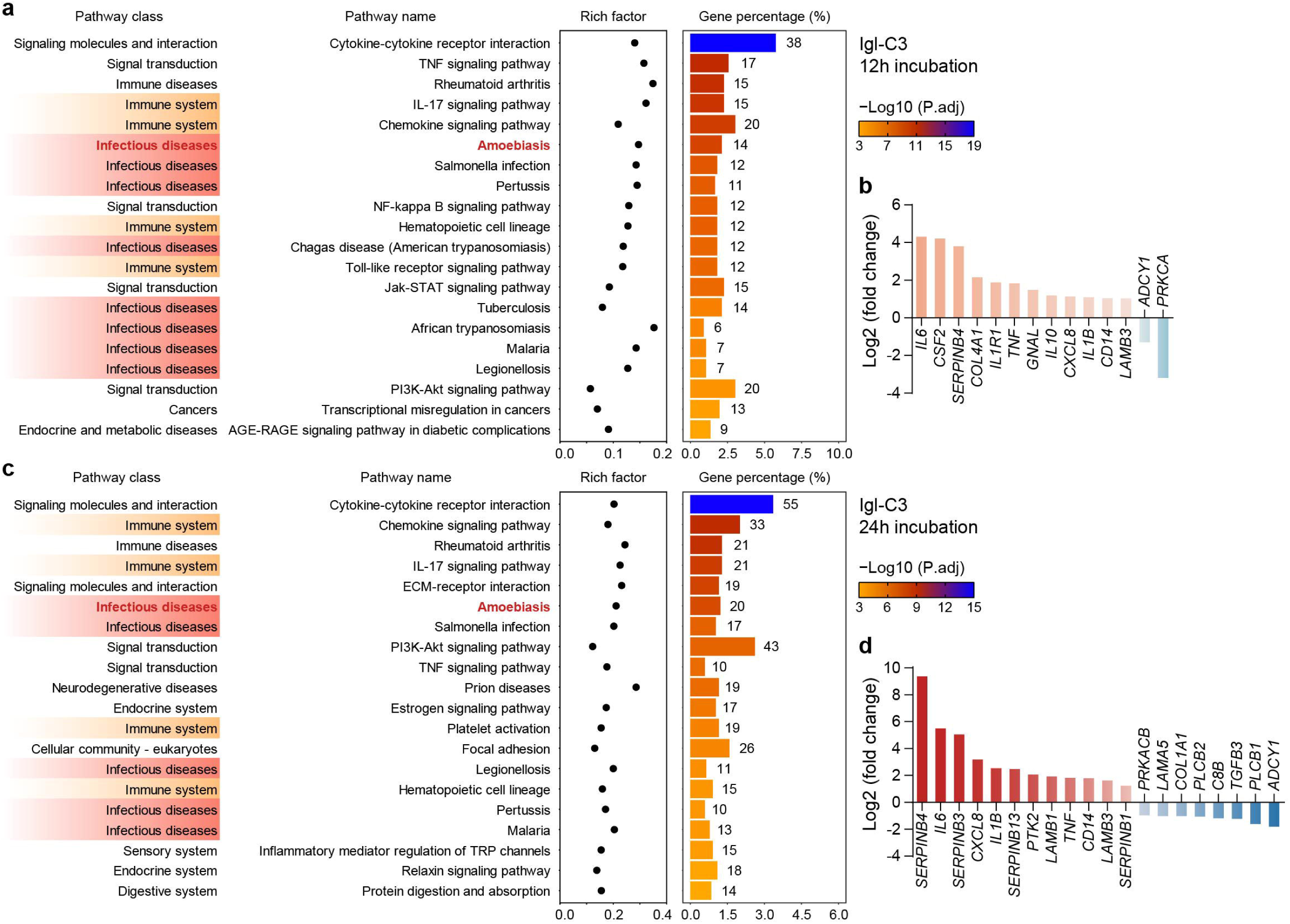

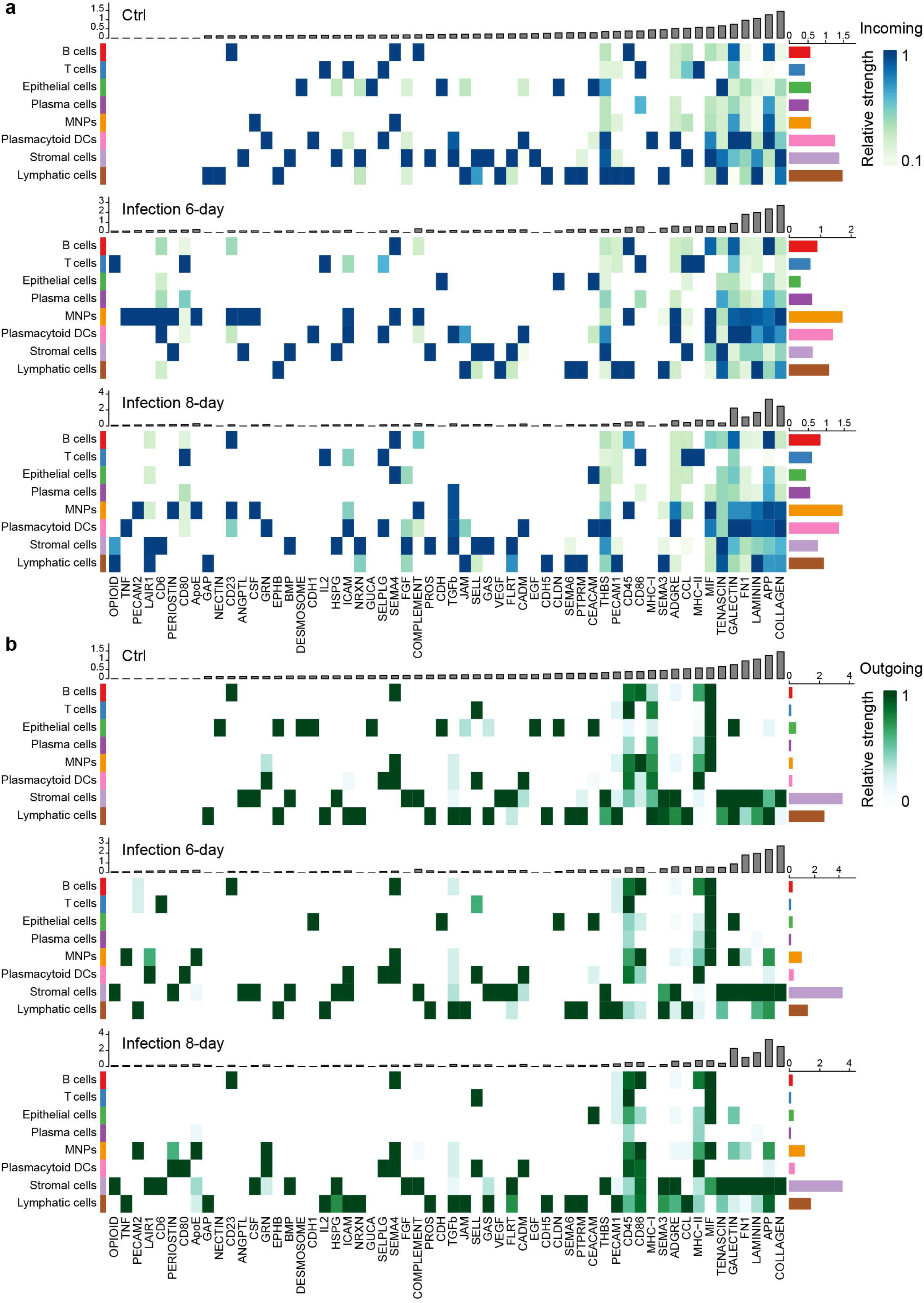

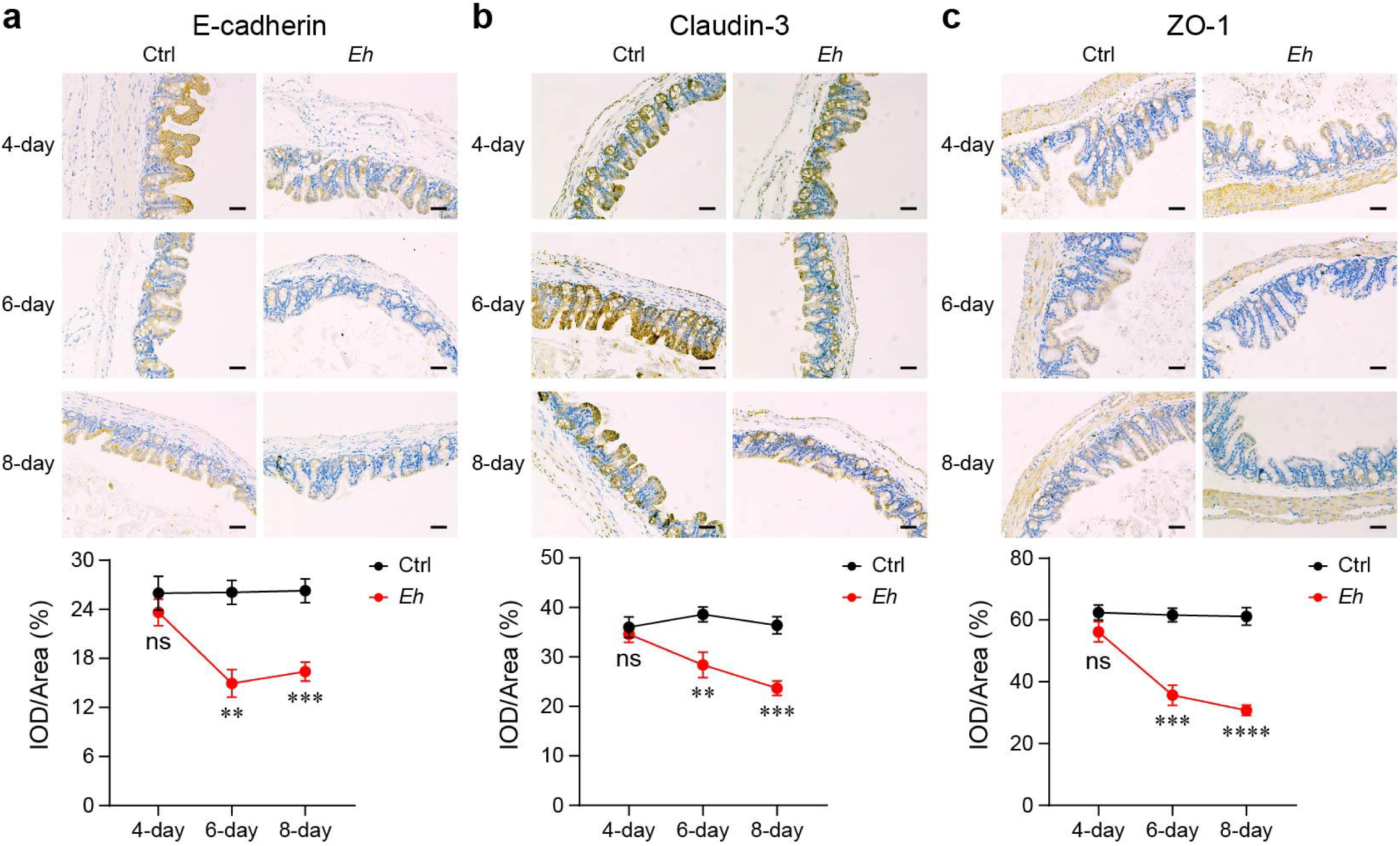

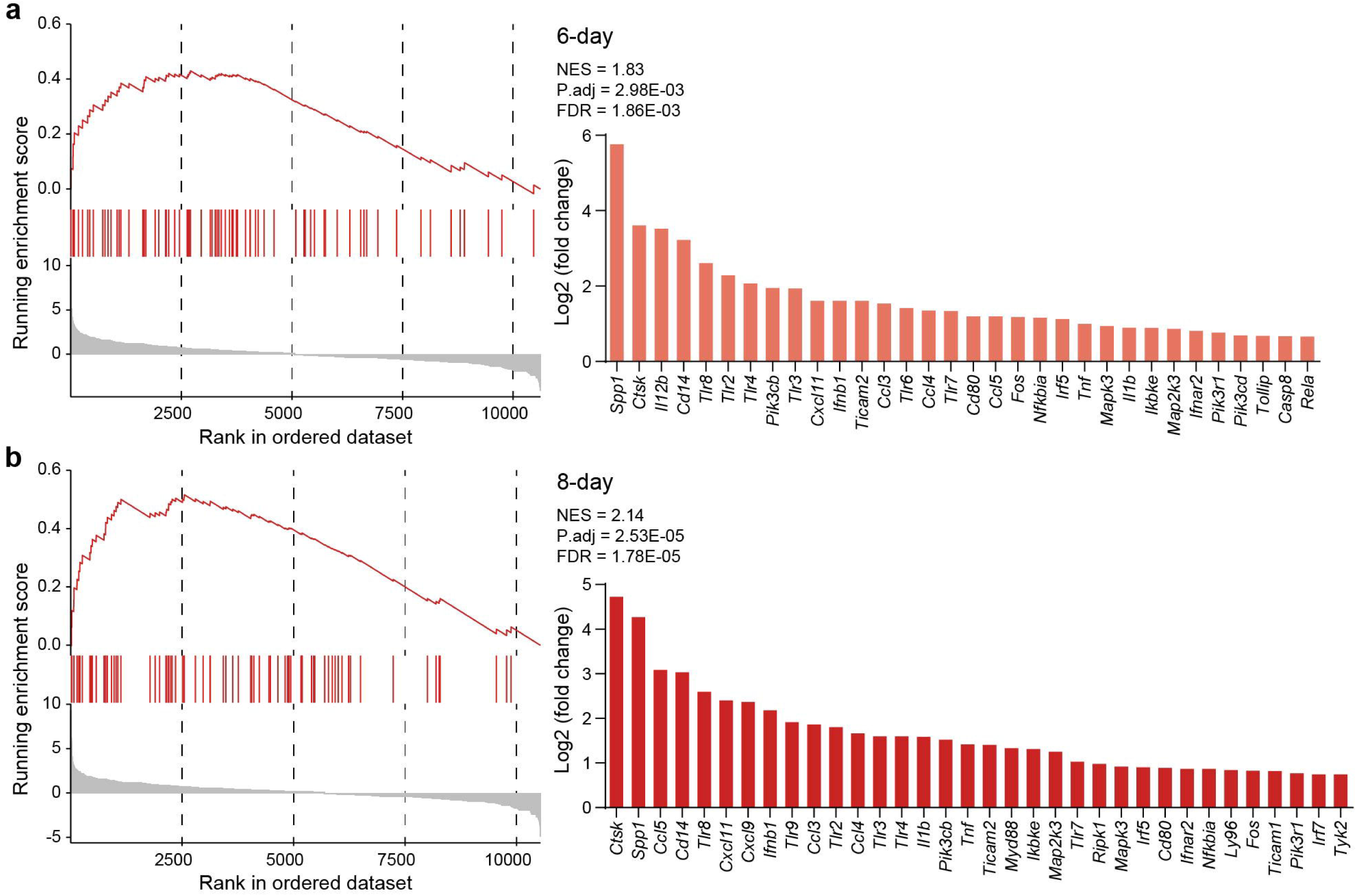

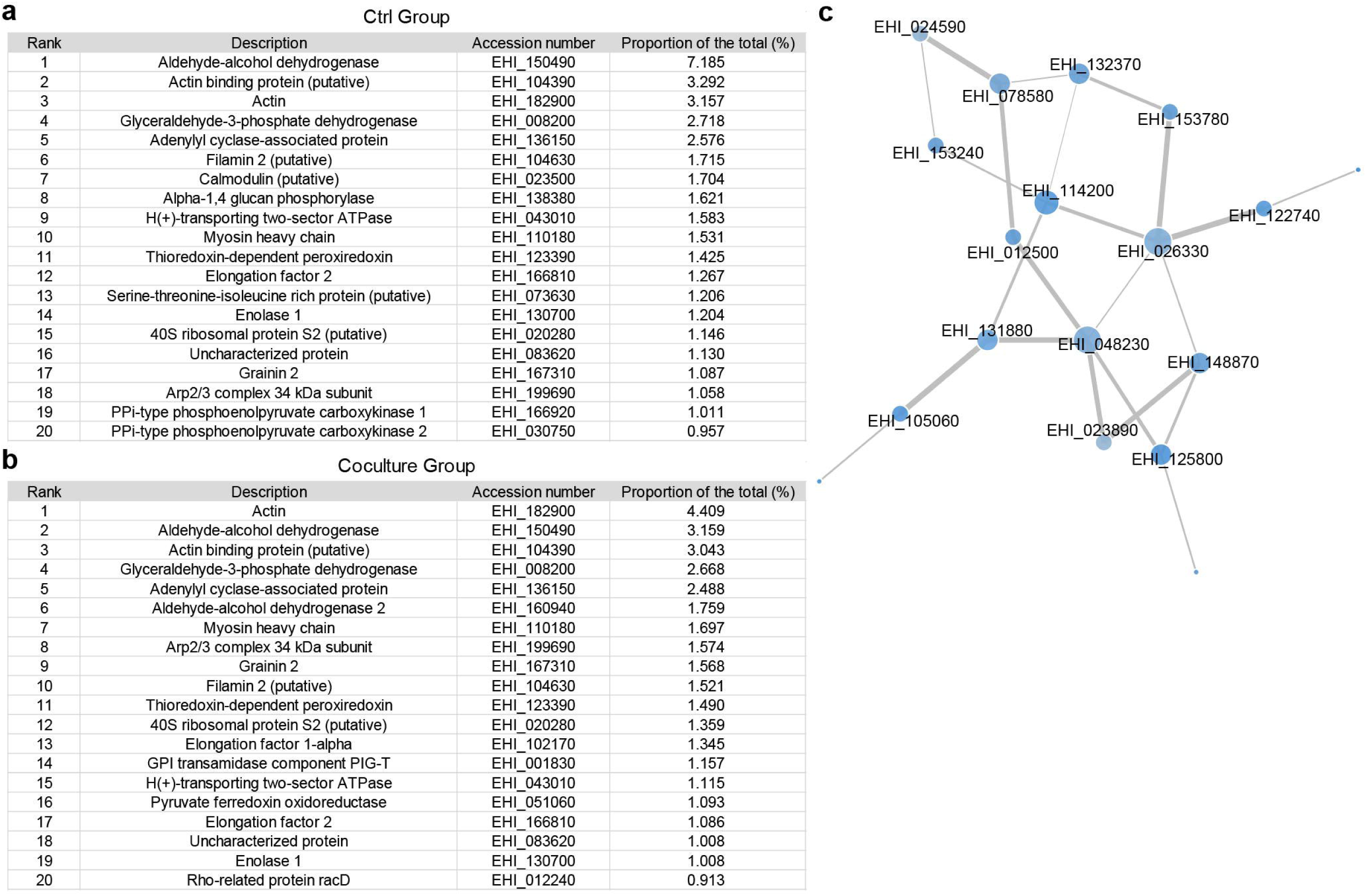

## References

1. Parfrey LW, Walters WA, Lauber CL, Clemente JC, Berg-Lyons D, Teiling C, et al. Communities of microbial eukaryotes in the mammalian gut within the context of environmental eukaryotic diversity. Frontiers in Microbiology 2014; 5.

2. Chabe M, Lokmer A, Segurel L. Gut Protozoa: Friends or Foes of the Human Gut Microbiota? Trends Parasitol 2017; 33:925–34.

3. Guillen N. Pathogenicity and virulence of Entamoeba histolytica, the agent of amoebiasis. Virulence 2023; 14:2158656.

4. Marie C, Petri WA, Jr. Regulation of virulence of Entamoeba histolytica. Annu Rev Microbiol 2014; 68:493–520.

5. Bansal D, Sehgal R, Chawla Y, Mahajan RC, Malla N. In vitro activity of antiamoebic drugs against clinical isolates of Entamoeba histolytica and Entamoeba dispar. Ann Clin Microbiol Antimicrob 2004; 3:27.

6. Chou A, Austin RL. Entamoeba histolytica Infection. StatPearls. Treasure Island (FL) ineligible companies. Disclosure: Richard Austin declares no relevant financial relationships with ineligible companies., 2025.

7. Manna D, Ehrenkaufer GM, Lozano-Amado D, Singh U. Entamoeba stage conversion: progress and new insights. Current Opinion in Microbiology 2020; 58:62–8.

8. Zhang H, Li Q, Zhou H, Feng M, Zhao Y, Zhou R, et al. Identification and characterization of a carbohydrate recognition domain-like region in Entamoeba histolytica Gal/GalNAc lectin intermediate subunit. Microbiol Spectr 2024; 12:e0053824.

9. Uddin MJ, Leslie JL, Petri WA, Jr. Host Protective Mechanisms to Intestinal Amebiasis. Trends Parasitol 2021; 37:165–75.

10. Groneberg M, Hoenow S, Marggraff C, Fehling H, Metwally NG, Hansen C, et al. HIF-1alpha modulates sex-specific Th17/Treg responses during hepatic amoebiasis. J Hepatol 2022; 76:160–73.

11. Chadee K, Meerovitch E. Entamoeba histolytica: early progressive pathology in the cecum of the gerbil (Meriones unguiculatus). Am J Trop Med Hyg 1985; 34:283–91.

12. Mortimer L, Chadee K. The immunopathogenesis of Entamoeba histolytica. Exp Parasitol 2010; 126:366–80.

13. Uribe-Querol E, Rosales C. Immune Response to the Enteric Parasite Entamoeba histolytica. Physiology (Bethesda) 2020; 35:244–60.

14. Prathap K, Gilman R. The histopathology of acute intestinal amebiasis. A rectal biopsy study. Am J Pathol 1970; 60:229–46.

15. Singh RS, Walia AK, Kanwar JR, Kennedy JF. Amoebiasis vaccine development: A snapshot on E. histolytica with emphasis on perspectives of Gal/GalNAc lectin. Int J Biol Macromol 2016; 91:258–68.

16. Ralston KS, Solga MD, Mackey-Lawrence NM, Somlata, Bhattacharya A, Petri WA, Jr. Trogocytosis by Entamoeba histolytica contributes to cell killing and tissue invasion. Nature 2014; 508:526–30.

17. Ravdin JI, Sperelakis N, Guerrant RL. Effect of ion channel inhibitors on the cytopathogenicity of Entamoeba histolytica. J Infect Dis 1982; 146:335–40.

18. Leippe M, Bruhn H, Hecht O, Grotzinger J. Ancient weapons: the three-dimensional structure of amoebapore A. Trends Parasitol 2005; 21:5–7.

19. Ralston KS, Petri WA, Jr. Tissue destruction and invasion by Entamoeba histolytica. Trends Parasitol 2011; 27:254–63.

20. Cornick S, Moreau F, Chadee K. Entamoeba histolytica Cysteine Proteinase 5 Evokes Mucin Exocytosis from Colonic Goblet Cells via alphavbeta3 Integrin. PLoS Pathog 2016; 12:e1005579.

21. Dey I, Keller K, Belley A, Chadee K. Identification and characterization of a cyclooxygenase-like enzyme from Entamoeba histolytica. Proc Natl Acad Sci U S A 2003; 100:13561–6.

22. Lauwaet T, Oliveira MJ, De Bruyne G, Bruchhaus I, Duchene M, Mareel M, et al. Entamoeba histolytica trophozoites transfer lipophosphopeptidoglycans to enteric cell layers. Int J Parasitol 2004; 34:549–56.

23. Gurung S, Perocheau D, Touramanidou L, Baruteau J. The exosome journey: from biogenesis to uptake and intracellular signalling. Cell Commun Signal 2021; 19:47.

24. Diaz-Godinez C, Rios-Valencia DG, Garcia-Aguirre S, Martinez-Calvillo S, Carrero JC. Immunomodulatory effect of extracellular vesicles from Entamoeba histolytica trophozoites: Regulation of NETs and respiratory burst during confrontation with human neutrophils. Front Cell Infect Microbiol 2022; 12:1018314.

25. Tsutsumi V, Shibayama M. Experimental amebiasis:: A selected review of some models. Arch Med Res 2006; 37:210–20.

26. Shimokawa C, Kabir M, Taniuchi M, Mondal D, Kobayashi S, Ali IKM, et al. Entamoeba moshkovskii is associated with diarrhea in infants and causes diarrhea and colitis in mice. Journal of Infectious Diseases 2012; 206:744–51.

27. Deloer S, Nakamura R, Mi-ichi F, Adachi K, Kobayashi S, Hamano S. Mouse models of amoebiasis and culture methods of amoeba. Parasitology International 2016; 65:520–5.

28. Penaranda C, Hung DT. Single-Cell RNA Sequencing to Understand Host-Pathogen Interactions. ACS Infect Dis 2019; 5:336–44.

29. Ho YT, Shimbo T, Wijaya E, Kitayama T, Takaki S, Ikegami K, et al. Longitudinal Single-Cell Transcriptomics Reveals a Role for Serpina3n-Mediated Resolution of Inflammation in a Mouse Colitis Model. Cell Mol Gastroenterol Hepatol 2021; 12:547–66.

30. Hong D, Kim HK, Yang W, Yoon C, Kim M, Yang CS, et al. Integrative analysis of single-cell RNA-seq and gut microbiome metabarcoding data elucidates macrophage dysfunction in mice with DSS-induced ulcerative colitis. Commun Biol 2024; 7:731.

31. Zhao Y, Li X, Zhou R, Zhang L, Chen L, Tachibana H, et al. Quantitative Proteomics Reveals Metabolic Reprogramming in Host Cells Induced by Trophozoites and Intermediate Subunit of Gal/GalNAc Lectins from Entamoeba histolytica. mSystems 2022; 7:e0135321.

32. Min XY, Feng M, Guan Y, Man SQ, Fu YF, Cheng XJ, et al. Evaluation of the C-Terminal Fragment of Gal/GalNAc Lectin Intermediate Subunit as a Vaccine Candidate against Amebic Liver Abscess. Plos Neglect Trop D 2016; 10.

33. Morbe UM, Jorgensen PB, Fenton TM, von Burg N, Riis LB, Spencer J, et al. Human gut-associated lymphoid tissues (GALT); diversity, structure, and function. Mucosal Immunol 2021; 14:793–802.

34. Cheng XJ, Hughes MA, Huston CD, Loftus B, Gilchrist CA, Lockhart LA, et al. Intermediate subunit of the Gal/GalNAc lectin of Entamoeba histolytica is a member of a gene family containing multiple CXXC sequence motifs. Infection and Immunity 2001; 69:5892–8.

35. Noll J, Helk E, Fehling H, Bernin H, Marggraff C, Jacobs T, et al. IL-23 prevents IL-13-dependent tissue repair associated with Ly6C(lo) monocytes in Entamoeba histolytica-induced liver damage. J Hepatol 2016; 64:1147–57.

36. Garcia-Hernandez V, Quiros M, Nusrat A. Intestinal epithelial claudins: expression and regulation in homeostasis and inflammation. Ann N Y Acad Sci 2017; 1397:66–79.

37. Ma T, Gu J, Zhao Y, Li S, Zou D, Ge D. EZH2-mediated suppression of CLDN1 leads to barrier dysfunction in PPI-refractory gastroesophageal reflux disease. Dig Liver Dis 2022; 54:776–83.

38. Hou D, Yu T, Lu X, Hong JY, Yang M, Zi Y, et al. Sirt2 inhibition improves gut epithelial barrier integrity and protects mice from colitis. Proc Natl Acad Sci U S A 2024; 121:e2319833121.

39. Su Y, Cheng SS, Ding YX, Wang LG, Sun MS, Man CX, et al. A comparison of study on intestinal barrier protection of polysaccharides from Hericium erinaceus before and after fermentation. International Journal of Biological Macromolecules 2023; 233.

40. Lozoya-Agullo I, Araujo F, Gonzalez-Alvarez I, Merino-Sanjuan M, Gonzalez-Alvarez M, Bermejo M, et al. Usefulness of Caco-2/HT29-MTX and Caco-2/HT29-MTX/Raji B Coculture Models To Predict Intestinal and Colonic Permeability Compared to Caco-2 Monoculture. Mol Pharm 2017; 14:1264–70.

41. Reale O, Huguet A, Fessard V. Co-culture model of Caco-2/HT29-MTX cells: A promising tool for investigation of phycotoxins toxicity on the intestinal barrier. Chemosphere 2020:128497.

42. Maldonado-Bernal C, Kirschning CJ, Rosenstein Y, Rocha LM, Rios-Sarabia N, Espinosa-Cantellano M, et al. The innate immune response to lipopeptidophosphoglycan is mediated by toll-like receptors 2 and 4. Parasite Immunology 2005; 27:127–37.

43. Li X, Feng M, Zhao Y, Zhang Y, Zhou R, Zhou H, et al. A Novel TLR4-Binding Domain of Peroxiredoxin From Entamoeba histolytica Triggers NLRP3 Inflammasome Activation in Macrophages. Front Immunol 2021; 12:758451.

44. Dai Y, Lu Q, Li P, Zhu J, Jiang J, Zhao T, et al. Xianglian Pill attenuates ulcerative colitis through TLR4/MyD88/NF-κB signaling pathway. J Ethnopharmacol 2023; 300.

45. Chen L, Fu W, Zheng L, Wang Y, Liang G. Recent progress in the discovery of myeloid differentiation 2 (MD2) modulators for inflammatory diseases. Drug Discov Today 2018; 23:1187–202.

46. Shi N, Li N, Duan XW, Niu HT. Interaction between the gut microbiome and mucosal immune system. Military Med Res 2017; 4.

47. Naylor C, Burgess S, Madan R, Buonomo E, Razzaq K, Ralston K, et al. Leptin receptor mutation results in defective neutrophil recruitment to the colon during Entamoeba histolytica infection. Mbio 2014; 5.

48. Diaz-Godinez C, Fonseca Z, Nequiz M, Laclette JP, Rosales C, Carrero JC. Entamoeba histolytica Trophozoites Induce a Rapid Non-classical NETosis Mechanism Independent of NOX2-Derived Reactive Oxygen Species and PAD4 Activity. Front Cell Infect Microbiol 2018; 8:184.

49. Sim S, Park SJ, Yong TS, Im KI, Shin MH. Involvement of beta 2-integrin in ROS-mediated neutrophil apoptosis induced by Entamoeba histolytica. Microbes Infect 2007; 9:1368–75.

50. Dickson-Gonzalez SM, de Uribe ML, Rodriguez-Morales AJ. Polymorphonuclear Neutrophil Infiltration Intensity as Consequence of Density in Amebic Colitis. Surg Infect 2009; 10:91-7.

51. Kammanadiminti SJ, Mann BJ, Dutil L, Chadee K. Regulation of Toll-like receptor-2 expression by the Gal-lectin of Entamoeba histolytica. FASEB J 2004; 18:155–7.

52. Ramos E, Olivos-Garcia A, Nequiz M, Saavedra E, Tello E, Saralegui A, et al. Entamoeba histolytica: apoptosis induced in vitro by nitric oxide species. Exp Parasitol 2007; 116:257–65.

53. Verkerke HP, Petri WA, Jr., Marie CS. The dynamic interdependence of amebiasis, innate immunity, and undernutrition. Semin Immunopathol 2012; 34:771–85.

54. Sellau J, Groneberg M, Fehling H, Thye T, Hoenow S, Marggraff C, et al. Androgens predispose males to monocyte-mediated immunopathology by inducing the expression of leukocyte recruitment factor CXCL1. Nature Communications 2020; 11.

55. Salata RA, Pearson RD, Ravdin JI. Interaction of human leukocytes and Entamoeba histolytica. Killing of virulent amebae by the activated macrophage. J Clin Invest 1985; 76:491–9.

56. Bagde S, Singh V. Effect of Prolonged Anti-HM1: IMSS Entamoeba histolytica Antibody Activity in Humoral and Cellular Immunity of Experimentally Induced Animal Model. Recent Pat Inflamm Allergy Drug Discov 2018; 12:87–95.

57. Newton K, Dixit VM. Signaling in innate immunity and inflammation. Cold Spring Harb Perspect Biol 2012; 4.

58. Tang DL, Kang R, Coyne CB, Zeh HJ, Lotze MT. PAMPs and DAMPs: signal 0s that spur autophagy and immunity. Immunol Rev 2012; 249:158–75.

59. Abed J, Emgard JE, Zamir G, Faroja M, Almogy G, Grenov A, et al. Fap2 Mediates Fusobacterium nucleatum Colorectal Adenocarcinoma Enrichment by Binding to Tumor-Expressed Gal-GalNAc. Cell Host Microbe 2016; 20:215–25.

60. Wu J, Li Q, Fu X. Fusobacterium nucleatum Contributes to the Carcinogenesis of Colorectal Cancer by Inducing Inflammation and Suppressing Host Immunity. Transl Oncol 2019; 12:846–51.

61. Zhang HZ, Jin K, Xiong KL, Jing WW, Pang Z, Feng M, et al. Disease-associated gut microbiome and critical metabolomic alterations in patients with colorectal cancer. Cancer Med-Us 2023; 12:15720–35.

62. Mian MF, Lauzon NM, Andrews DW, Lichty BD, Ashkar AA. FimH can directly activate human and murine natural killer cells via TLR4. Mol Ther 2010; 18:1379–88.

63. Jeisy-Scott V, Kim JH, Davis WG, Cao WP, Katz JM, Sambhara S. TLR7 Recognition Is Dispensable for Influenza Virus A Infection but Important for the Induction of Hemagglutinin-Specific Antibodies in Response to the 2009 Pandemic Split Vaccine in Mice. J Virol 2012; 86:10988–98.

64. Goff PH, Hayashi T, Martnez-Gil L, Corr M, Crain B, Yao SY, et al. Synthetic Toll-Like Receptor 4 (TLR4) and TLR7 Ligands as Influenza Virus Vaccine Adjuvants Induce Rapid, Sustained, and Broadly Protective Responses. J Virol 2015; 89:3221–35.

65. Ivory CP, Chadee K. Activation of dendritic cells by the Gal-lectin of Entamoeba histolytica drives Th1 responses in vitro and in vivo. Eur J Immunol 2007; 37:385–94.

66. Ivory CP, Prystajecky M, Jobin C, Chadee K. Toll-like receptor 9-dependent macrophage activation by Entamoeba histolytica DNA. Infect Immun 2008; 76:289–97.

67. Spalinger MR, Sayoc-Becerra A, Santos AN, Shawki A, Canale V, Krishnan M, et al. PTPN2 Regulates Interactions Between Macrophages and Intestinal Epithelial Cells to Promote Intestinal Barrier Function. Gastroenterology 2020; 159:1763-+.

68. Chen GX, Ran X, Li B, Li YH, He DW, Huang BX, et al. Sodium Butyrate Inhibits Inflammation and Maintains Epithelium Barrier Integrity in a TNBS-induced Inflammatory Bowel Disease Mice Model. Ebiomedicine 2018; 30:317–25.

69. Bain CC, Scott CL, Uronen-Hansson H, Gudjonsson S, Jansson O, Grip O, et al. Resident and pro-inflammatory macrophages in the colon represent alternative context-dependent fates of the same Ly6C monocyte precursors. Mucosal Immunology 2013; 6:498–510.

70. Sewell GW, Rahman FZ, Levine AP, Jostins L, Smith PJ, Walker AP, et al. Defective tumor necrosis factor release from Crohn’s disease macrophages in response to Toll-like receptor activation: relationship to phenotype and genome-wide association susceptibility loci. Inflamm Bowel Dis 2012; 18:2120–7.

71. Torretta S, Scagliola A, Ricci L, Mainini F, Di Marco S, Cuccovillo I, et al. D-mannose suppresses macrophage IL-1beta production. Nat Commun 2020; 11:6343.

72. Alhendi A, Naser SA. The dual role of interleukin-6 in Crohn’s disease pathophysiology. Front Immunol 2023; 14:1295230.

73. Wojcik GL, Marie C, Abhyankar MM, Yoshida N, Watanabe K, Mentzer AJ, et al. Genome-Wide Association Study Reveals Genetic Link between Diarrhea-Associated Infection and Inflammatory Bowel Disease. Mbio 2018; 9.

74. Panarelli NC. Infectious Mimics of Inflammatory Bowel Disease. Modern Pathol 2023; 36.

75. Zhang L, Cheng D, Zhang J, Tang H, Li F, Peng Y, et al. Role of macrophage AHR/TLR4/STAT3 signaling axis in the colitis induced by non-canonical AHR ligand aflatoxin B1. J Hazard Mater 2023; 452:131262.

76. Xie Y, Irwin S, Nelson B, van Daelen M, Fontenot L, Jacobs JP, et al. Citrulline Inhibits Clostridioides difficile Infection With Anti-inflammatory Effects. Cell Mol Gastroenterol Hepatol 2025; 19:101474.

77. Wu ZY, Wang LL, Li JY, Wang LF, Wu ZD, Sun X. Extracellular Vesicle-Mediated Communication Within Host-Parasite Interactions. Frontiers in Immunology 2019; 9.

78. Ujang JA, Kwan SH, Ismail MN, Lim BH, Noordin R, Othman N. Proteome analysis of excretory-secretory proteins of HM1:IMSS via LC-ESI-MS/MS and LC-MALDI-TOF/TOF. Clin Proteom 2016; 13.

79. Nievas YR, Lizarraga A, Salas N, Cóceres VM, de Miguel N. Extracellular vesicles released by anaerobic protozoan parasites: Current situation. Cellular Microbiology 2020; 22.

80. Sharma M, Morgado P, Zhang HB, Ehrenkaufer G, Manna D, Singh U. Characterization of Extracellular Vesicles from Identifies Roles in Intercellular Communication That Regulates Parasite Growth and Development. Infection and Immunity 2020; 88.

81. Honecker B, Barreiter VA, Hohn K, Horvath B, Harant K, Metwally NG, et al. Entamoeba histolytica extracellular vesicles drive pro-inflammatory monocyte signaling. PLoS Negl Trop Dis 2025; 19:e0012997.

82. Chowdhury D, Sharma M, Jahng JWS, Singh U. Extracellular Vesicles Derived From Entamoeba histolytica Have an Immunomodulatory Effect on THP-1 Macrophages. Journal of Parasitology Research 2024; 2024.

83. Rahim Z, Raymonddenise A, Sansonetti P, Guillen N. Localization of Myosin Heavy Chain-a in the Human Pathogen Entamoeba-Histolytica. Infection and Immunity 1993; 61:1048–54.

84. Arhets P, Gounon P, Sansonetti P, Guillen N. Myosin-Ii Is Involved in Capping and Uroid Formation in the Human Pathogen Entamoeba-Histolytica. Infection and Immunity 1995; 63:4358–67.

85. Leroy A, Debruyne G, Mareel M, Nokkaew C, Bailey G, Nelis H. Contact-Dependent Transfer of the Galactose-Specific Lectin of Entamoeba-Histolytica to the Lateral Surface of Enterocytes in Culture. Infection and Immunity 1995; 63:4253–60.

86. Ng YL, Olivos-García A, Lim TK, Noordin R, Lin QS, Othman N. Entamoeba histolytica: Quantitative Proteomics Analysis Reveals Putative Virulence-Associated Differentially Abundant Membrane Proteins. American Journal of Tropical Medicine and Hygiene 2018; 99:1518–29.

87. Lozano-Mendoza J, Ramírez-Montiel F, Rangel-Serrano A, Páramo-Pérez I, Mendoza-Macías CL, Saavedra-Salazar F, et al. Attenuation of In Vitro and In Vivo Virulence Is Associated with Repression of Gene Expression of AIG1 Gene in Entamoeba histolytica. Pathogens 2023; 12.

88. Leon-Coria A, Kumar M, Moreau F, Chadee K. Defining cooperative roles for colonic microbiota and Muc2 mucin in mediating innate host defense against Entamoeba histolytica. Plos Pathogens 2018; 14.

