## Supplemental Methods for "*Entamoeba histolytica* Gal/GalNAc lectin intermediate subunit promotes inflammation and epithelial damage in intestinal amebiasis through its C3 region"

**E****xpression and purification of different Igl-C3 proteins**

*Entamoeba histolytica* Igl has two subtypes, Igl1 and Igl2, whose C3 regions are 99% identical and 100% positive ^1^. In the present study, since amoebic Igl mostly exists in the form of Igl1 subtype, the C3 segment of the *Igl1* gene (GenBank: AF337950.1) was amplified via PCR to construct different vectors. Total cDNA from *E. histolytica* HM-1:IMSS trophozoites was used as the template.

For eukaryotic Igl-C3 expression, the FLAG-linker-Igl-C3 sequence was subcloned into a pCMV6 plasmid (#PS100001; OriGene) via double digestion of restriction enzymes Sgf I and Mlu I, and the pCMV6-Igl-C3 plasmid was then amplified within *Escherichia coli* DH5α competent cells and extracted with an EndoFree Plasmid Maxi Kit (Qiagen, Hilden, Germany). Plasmids were transiently transfected into HEK 293 cells using polyethyleneimine (PEI) for protein expression. After 48 h of shaking culture at 37°C, cells were harvested and lysed using 25 mM Tris-HCl pH 7.4 with 150 mM NaCl, 1% (v/v) NP-40, and 1 mM EDTA. The eukaryotic protein was purified by anti-FLAG tag affinity chromatography, eluted with 0.1 M glycine (pH 2.75), then confirmed using sodium dodecyl sulfate-polyacrylamide gel electrophoresis (SDS-PAGE).

Prokaryotic Igl-C3 was only used for protein affinity detection, whose expression and purification were performed as previously described ^1^. Briefly, the Igl-C3 sequence was amplified and subcloned into a pET-19b plasmid (Novagen, Darmstadt, Germany) at the XhoI restriction site. After being transformed with the subcloned plasmid, *E. coli* BL21 Star(DE3)pLysS competent cells (Weidibio, Shanghai, China) were cultured in Luria-Bertani medium containing 100 µg/mL ampicillin, then induced with 1 mM isopropyl-β-d-thiogalactopyranoside (Amresco, Solon, OH, USA). The bacteria were incubated at 37°C for 3 h, then harvested and sonicated. The supernatants were filtrated via a 0.45-µm membrane, further purified using a Ni-NTA (nitrilotriacetic acid) His-Bind resin kit (Novagen, Darmstadt, Germany). A Toxin Sensor Gel Clot Endotoxin Assay Kit (Genscript, Nanjing, China) was applied to detect endotoxin level in the prokaryotic protein, which conformed to the national standard of the People's Republic of China for medical products (GB/t14233.2–2005). All corresponding primers are listed in Table S2.

**Purification of** **native Igl protein**

For the purification of native Igl protein from *E. histolytica* HM-1:IMSS trophozoites, the anti-Igl monoclonal antibody (mAb) EH3015 was first conjugated to CNBr-activated Sepharose 4B gel ^2^. *E. histolytica* trophozoites in the logarithmic growth phase were harvested and suspended in 50 mM Tris-HCl buffer (pH 8.3) containing 150 mM NaCl, 0.5% (v/v) Nonidet P-40, and 5 mM EDTA disodium salt prior to sonication. Subsequently, the soluble fraction was loaded onto an mAb EH3015-bound affinity column. After extensive washing, native Igl protein was eluted with 0.2 N acetic acid (pH 3.0) and dialyzed immediately in 20 mM Tris-HCl buffer (pH 8.0). The purity of protein samples was confirmed using SDS-PAGE.

**Quantitative real-time RT-PCR**

Quantitative real-time RT-PCR was implemented to detect gene expressions in the host cells and *E. histolytica* trophozoites. Total RNA from the cells or trophozoites was purified with a RNeasy Plus Mini Kit (Qiagen, Dusseldorf, Germany), then reverse transcribed using a PrimeScript 1st strand cDNA Synthesis Kit (TaKaRa, Shiga, Japan). In accordance with manufacturer's protocols, qRT-PCR was conducted by a SYBR Premix Ex Taq (TaKaRa, Shiga, Japan) in a final reaction volume of 20 µL on an ABI 7500 real-time PCR system (Applied Biosystems, CA, USA). Human *GAPDH*, *TNF*, *IL1B*, *IL6*, *IL10*, *IL23A*, *TGFB1*, *CDH1*, *CLDN3*, *TJP1*, *MYD88*, *LY96* genes, mouse *Actb*, *Tnf*, *Il1b*, *Il6*, *Nos2*, *Arg1* genes, and *E. histolytica* *Actin*, *Igl1*, *Igl2* genes were set as detection targets. Reactions were performed in 96-well plates under the following amplification cycling conditions: 30 s at 95°C, 40 cycles of 5 s at 95°C, and 35 s at 60°C. Gene expressions were analyzed using the 2^−ΔΔCt^ method. The primer sets are shown in Table S3.

**Western blotting**

Western blotting was implemented to detect the protein expressions in host cells and *E. histolytica* extracellular vesicles (EVs). Protein samples were pelleted by centrifugation, separated on gradient polyacrylamide gels, then electrotransferred onto polyvinylidene difluoride membranes (General Electric Co., Schenectady, NY, USA). After blocking with 5% skim milk in tris buffered saline (TBS), membranes were incubated with primary antibodies targeting human β-Actin, TNF-α, IL-1β, IL-6, MyD88, NF-κB p65, phospho-NF-κB p65, MD2 (1:1000), and *E. histolytica* Igl (1:500) overnight at 4℃. Subsequently, the membranes were incubated with horseradish peroxidase (HRP)-conjugated goat anti-mouse/rabbit/Syrian hamster IgG H&L (1:5000) at room temperature for 1 h. The proteins were finally detected using an enhanced chemiluminescence (ECL) Western Blotting Substrate Kit (Tanon, Shanghai, China). Information regarding the antibodies used is detailed in Table S4.

**L****aser confocal microscopy**

To detect the expression and distribution of phospho-NF-κB protein in U937 cells, laser confocal microscopy was implemented using glass-bottom dishes (D35C4-20-1.5-N; Cellvis, Mountain View, CA, USA). The cells were fixed with 4% paraformaldehyde, permeabilized with 0.2% Triton X-100, then blocked with 3% BSA in phosphate buffered saline (PBS). Phospho-NF-κB rabbit mAb (1:1,000, CST) was used as the primary antibody, while Alexa Fluor 594 goat anti-rabbit IgG H&L (1:500, Thermo Fisher) was used as the secondary antibody. Before observing under a laser confocal microscope (SP8; Leica Microsystems, Wetzlar, Germany), dishes were stained with 0.25 mg/mL DAPI (1:200) for nuclear visualization.

**Mucus staining of the epithelial cell monolayer**

To compare the simulation effect of coculture with different proportions of Caco-2 and HT29-MTX-E12 cells (3:1 or 9:1 ratio) on the human intestinal epithelium, alcian blue and eosin staining was performed on the cell monolayer. First, transwell inserts with Caco-2/HT29-MTX-E12 cocultured cells were washed twice with PBS, and fixed with 4% paraformaldehyde in both the apical and basal compartments for 20 min at room temperature. After washing with distilled water, 1% alcian blue 8GX in 3% acetic acid solution (66011; Sigma-Aldrich, St. Louis, MO, USA) was added to the apical compartment for 30 min to visualize acidic mucins. Subsequently, the cells were washed and counterstained with 1% eosin solution for 1 min (also added to the apical side). The polyester membranes were then removed using a scalpel blade, and embedded in an OCT-embedding matrix. Frozen embedded polyester membranes were finally cut into 5 μm vertical cross sections using a Leica CM1950 cryotome (Leica, Wetzlar, Germany).

**TEER and transepithelial flux assays**

For the three-dimensional triple culture of human Caco-2, HT29-MTX-E12, and U937 cells, the two intestinal epithelial cell lines (3:1 ratio; 8 × 10^4^ cells in total) were first seeded on top of the transwell inserts (24-well format) and grown for mucus production. Later, the inserts were transferred onto PMA-differentiated U937 cells (8 × 10^5^ cells) seeded in 24-well culture plates. As a comparison, Igl-C3 protein (2 μg/mL) was added to the intestinal epithelial cells in the presence or absence of U937 cells.

In accordance with manufacturer's protocols, the integrity of the epithelial cell monolayer was determined by measuring transepithelial electronic resistance (TEER) with a Millicell ERS 3.0 Digital Voltohmmeter (MERS03000; Merck Millipore, Bedford, MA, USA). TEER value was calculated by the following formula: TEER (Ω·cm^2^) = TEER (Ω) × effective membrane area (cm^2^).

Epithelial cell monolayer permeability was further evaluated through fluorescein isothiocyanate (FITC)-dextran (FD4; Sigma-Aldrich, St. Louis, MO, USA) transmission. FITC-dextran (1 mg/mL) was applied to the apical side of transwell inserts, then incubated for 2 h at 37℃. Solutions on the basal side were collected, and the fluorescence intensities of the transferred FITC-dextran (490-nm excitation; 525-nm emission) were finally measured using a BioTek Synergy H1 Microplate Reader (Agilent BioTek, Vermont, USA).

**Detection of cytokine secretion and protein affinity by ELISA**

After collecting the supernatant of human U937 cells, enzyme-linked immunosorbent assay (ELISA) kits were used to detect the levels of TNF-α (DL-TNFa-Hu; DLdevelop, Wuxi, China), IL-1β (DL-IL1b-Hu; DLdevelop), IL-6 (DL-IL6-Hu; DLdevelop), IL-10 (DL-IL10-Hu; DLdevelop), IL-23 (DL-IL23-Hu; DLdevelop), and TGF-β1 (DL-TGFb1-Hu; DLdevelop) in accordance with manufacturer's protocols.

ELISA was also implemented to analyze the affinities of human MD2 protein (Ab238343; Abcam, Cambridge, United Kingdom) to eukaryotic Igl-C3, prokaryotic Igl-C3, and LPS (L2880; Sigma-Aldrich). After coating with Igl-C3 or LPS (2 μg), ELISA microplates (92592; Corning, Kennebunk, NY, USA) were blocked with PBS containing 1% skim milk. The MD2 protein (2 μg) was added for overnight incubation at 4℃. MD2 rabbit polyclonal antibody (1:200, Proteintech) and HRP-conjugated goat anti-rabbit IgG H&L (1:500; Abcam) were used as primary and secondary antibodies, respectively. With a TMB-Elisa liquid substrate solution (GPC, Beijing, China), optical density at 450 nm was measured upon incubation at room temperature for 10 min.

**Detection of** **p****rotein affinity via surface plasmon resonance**

According to the manufacturer's instructions, the functional affinities of eukaryotic Igl-C3/human MD2 (Ab238343; Abcam), eukaryotic Igl-C3/human TLR4 (Ab233665; Abcam), and native Igl/human MD2 were measured by the biosensor-based surface plasmon resonance (SPR) technique using an automatic apparatus BIAcore T200 (GE Healthcare Life Sciences, USA). Eukaryotic Igl-C3 or purified native Igl protein from *E. histolytica* was coupled to a chip (GE Healthcare Life Sciences), then serial dilutions of human MD2 or TLR4 were loaded to detect the binding responses. K_D_ values were calculated using Biacore analysis software.

**Inhibitor and siRNA treatments**

Inhibitor and small interfering RNA (siRNA) treatments were performed on human U937 cells. Before each treatment, U937 cells were first seeded in 24-well culture plates (for qPCR), 6-well culture plates (for western blotting), or glass-bottom dishes (for laser confocal microscopy), then pretreated with PMA for 24 h for differentiation. Subsequently, 1 μM TLR1/2 inhibitor CU-CPT22 (HY-108471; Merck Millipore, Darmstadt, Germany), 1 μM TLR4 inhibitor TAK-242 (A3850; APExBIO, Huston, USA), or 5 μM MD2 inhibitor MD2-IN-1 (HY-103483; Merck Millipore, Darmstadt, Germany) was added. After 2 h incubation, the medium was changed and the cells were stimulated with 2 μg/mL Igl-C3 protein for another 12 or 24 h.

According to the manufacturer's instructions, Lipofectamine 2000 Transfection Reagent (11668-019; Invitrogen, Carlsbad, CA, USA) was used for siRNA transfection (40 nM) to silence the *GAPDH*, *MYD88*, and *LY96* genes. The medium was changed after 6 h, and 2 μg/mL Igl-C3 protein was added for stimulation after 24 h. The corresponding oligonucleotides are listed in Table S5.

**EV isolation**

For amoebic EV isolation, 2 × 10^6^ *E. histolytica* trophozoites were first seeded into T25 culture flasks under standard growth conditions. After parasite attachment, the flask was washed twice with 37℃ 2% glucose-PBS, then 5 mL serum-free YIMDHA-S medium was added, which contained 1 × 10^7^ differentiated U937 cells (1:5 ratio) or not. Parasites and human cells were incubated anaerobically for 3 h at 37°C, then the culture supernatant was collected carefully, centrifuged at 500 × *g* for 5 min, and filtered through 0.22 μm filters to remove cell debris.

The EVs were purified with a Total Exosome Isolation Reagent (4478359; Invitrogen, Carlsbad, CA, USA) according to the manufacturer's protocol. After isolation reagent addition, the mixture was incubated overnight at 4°C, then centrifuged at 10,000 × *g* for 1 h to pellet the EVs. Finally, EV pellets were resuspended in PBS for subsequent analyses. SDS-PAGE and Coomassie Brilliant Blue staining were used to identify the protein components in EVs.

**Single-cell RNA-seq: Sample preparation and library construction**

At 6 and 8 d post-inoculation with *E. histolytica* trophozoites, C3H/HeNCrl mice were euthanized and their fresh cecal tissues were collected for single-cell isolation. After removing the residual contents, the mouse ceca were washed thrice with cold PBS and cut into small pieces with a diameter of roughly 5 mm. According to the manufacturer's instructions, cecal pieces were digested with a PythoN Tissue Dissociator (SGR-TDAp101; Singleron, Suzhou, China) and supporting enzymes. Subsequently, tissues were filtered through 30 μm MACS SmartStrainers (130-098-458; Miltenyi Biotec, Auburn, CA, USA), then pelleted by centrifugation at 4°C and washed with cold DMEM medium to remove the debris. Upon confirming the cell viability with ReadyCount Green/Red Stains (A49905; Invitrogen, Carlsbad, CA, USA), concentration of each sample was adjusted to 1,000 cells/μL.

Single cells of mouse ceca were loaded onto a 10X Genomics Chromium iX instrument, then single-cell libraries were prepared using a Single Cell 3′ library and Gel Bead Kit v3.1 in accordance with the manufacturer’s protocols. Generation of gel beads in emulsions (GEMs), barcoding, GEM-RT clean-up, complementary DNA (cDNA) amplification, and library construction were successively performed. After assessing cDNA quality with a Qubit dsDNA HS Assay Kit (Invitrogen, Carlsbad, CA, USA), the final library pool was sequenced on an Illumina NovaSeq 6000 platform using 150 base-pair paired-end reads.

**Single-cell RNA-seq: Data processing and cell type annotation**

Sample demultiplexing, barcode processing, alignment to the mouse genome GRCm39, and raw gene expression counting were performed using the Cell Ranger version 8.0.0 pipeline ^3^. Ambient RNA was removed by the CellBender version 0.3.2. Using R version 4.4.1, raw single-cell expression matrices were then analyzed by the Seurat package version 5.2.1. The criteria for removing low-quality cells were set as follows: (1) > 5% UMIs derived from the mitochondrial genome; (2) < 200 genes or >10,000 genes; (3) <500 UMIs. Doublets were identified by the DoubletFinder package version 2.0.3 with default settings, leaving 68,058 cells (from the ceca of nine different mice) and 23,208 genes for further analysis. After gene expression normalization using SCTransform version 2.0.3.5, single-cell RNA-seq data matrices from different samples were integrated using the FindIntegrationAnchors and IntegrateData functions to remove batch effects. Subsequently, linear dimensional reduction was performed with the RunPCA function, nonlinear dimensional reduction was performed with the RunUMAP function, and cluster analysis was performed with the FindNeighbors and FindClusters functions.

For cell type annotation and cluster marker identification, the shared nearest neighbor (SNN) clustering algorithm in Seurat was used, with a resolution of 1.6. Cell types were assigned by examining the expression of marker genes and the top differentially expressed genes (DEGs) in each cluster ^4^. The cell types were visualized using the Uniform Manifold Approximation and Projection (UMAP) clustering algorithm. Signature genes of each cell cluster were identified by the FindAllMarkers function in Seurat. Percentages of the different cell types were calculated accordingly.

**Single-cell RNA-seq: DEG identification and functional enrichment**

DEGs were identified by the FindMarkers function in Seurat with the following parameters: min.pct = 0.1; test.use = bimod. Genes with an average log2 fold change (log2FC) value > 0.585 and adjusted p value < 0.05 were identified as upregulated DEGs, while genes with an average log2FC value < -0.585 and adjusted p value < 0.05 were identified as downregulated DEGs. Using the ClusterProfiler package version 4.12.2, Gene Ontology (GO) and Kyoto Encyclopedia of Genes and Genomes (KEGG) enrichment analyses were further conducted with gene set enrichment analysis (GSEA). Cell-cell communication analysis was performed using the CellChat package version 2.1.2. Finally, in the bioinformatics section, R version 4.4.1 was employed for all statistical analyses, with an adjusted p value < 0.05 being considered statistically significant.

**T****ranscriptomic sequencing and subsequent analysis of human U937 cells**

Differentiated U937 cells were stimulated with 2 μg/mL Igl-C3 for 12 or 24 h, then the total RNA was extracted using a RNeasy Plus Mini Kit (Qiagen). Each sample consisted of three independent biological replicates. After evaluating nucleic acid quality via gel electrophoresis and a Qubit 4 Fluorometer (Q33226; Thermo Scientific, Waltham, MA, USA), 200 ng of total RNA was used for strand-specific sequencing library construction with a TruSeq RNA Sample Preparation Kit (Illumina, San Diego, CA, USA). Sequencing was performed using an Illumina NovaSeq 6000 instrument.

Raw data were processed using Skewer version 0.2.2, data quality was assessed using FastQC version 0.11.5, and clean reads were aligned to the human genome using STAR 2.5.3a. Setting |log2FC| value > 1 and adjusted p value < 0.05 as the criteria, DEGs between multiple experimental and control groups were identified by DESeq2 version 1.16.1. Using R version 4.4.1, KEGG and GO enrichment analyses were further conducted based on the DEGs. Enriched pathways and terms with adjusted p value < 0.05 were considered significant.

**Q****uantitative** **proteomics and subsequent analysis of amoebic EVs**

Four-dimensional label-free quantitative proteomics was applied to analyze changes in protein composition in amoebic EVs after co-incubation of *E. histolytica* trophozoites with human macrophages. Each sample consisted of three independent biological replicates. Upon protein extraction, concentration determination, and trypsin digestion, the peptides were desalted on a Sep-Pak column (Waters, Framingham, MA, USA) according to the manufacturer's instructions. Subsequently, the peptides were separated using a NanoElute Ultra Performance Liquid Chromatography system (Bruker Daltonics, Billerica, MA, USA) for liquid chromatography-tandem mass spectrometry (LC-MS) analysis. MS data were processed using PEAKS Online 1.5 (Bioinformatics Solutions Inc., Waterloo, Canada), and the data library was downloaded from the National Center for Biotechnology Information (NCBI) database. To screen for differential proteins, |log2FC| value > 1 and adjusted p value < 0.05 were set as the criteria. GO enrichment analysis was further conducted using R version 4.4.1, and enriched terms with an adjusted p value < 0.05 were considered significant.

**References**

1.	Zhang H, Li Q, Zhou H, Feng M, Zhao Y, Zhou R, et al. Identification and characterization of a carbohydrate recognition domain-like region in Entamoeba histolytica Gal/GalNAc lectin intermediate subunit. Microbiol Spectr 2024; 12:e0053824.

2.	Zhao Y, Li X, Zhou R, Zhang L, Chen L, Tachibana H, et al. Quantitative Proteomics Reveals Metabolic Reprogramming in Host Cells Induced by Trophozoites and Intermediate Subunit of Gal/GalNAc Lectins from Entamoeba histolytica. mSystems 2022; 7:e0135321.

3.	Luecken MD, Theis FJ. Current best practices in single-cell RNA-seq analysis: a tutorial. Mol Syst Biol 2019; 15.

4.	Ho YT, Shimbo T, Wijaya E, Kitayama T, Takaki S, Ikegami K, et al. Longitudinal Single-Cell Transcriptomics Reveals a Role for Serpina3n-Mediated Resolution of Inflammation in a Mouse Colitis Model. Cell Mol Gastroenterol Hepatol 2021; 12:547-66.
