## Supplemental Tables for "*Entamoeba histolytica* Gal/GalNAc lectin intermediate subunit promotes inflammation and epithelial damage in intestinal amebiasis through its C3 region"

**Table S1.** **Gene expression for each cell type in single-cell RNA-seq.**

| **Cell type** | **Upregulated** | **Downregulated** | **Not changed** |
| --- | --- | --- | --- |
| **B cells** | | | |
| Infection 6-day vs Ctrl | 121 | 26 | 7,568 |
| Infection 8-day vs Ctrl | 86 | 124 | 6,805 |
| **Epithelial cells** | | | |
| Infection 6-day vs Ctrl | 60 | 366 | 10,561 |
| Infection 8-day vs Ctrl | 349 | 565 | 10,217 |
| **Lymphatic cells** | | | |
| Infection 6-day vs Ctrl | 0 | 0 | 7,779 |
| Infection 8-day vs Ctrl | 0 | 0 | 10,167 |
| **MNPs** | | | |
| Infection 6-day vs Ctrl | 236 | 85 | 10,845 |
| Infection 8-day vs Ctrl | 152 | 31 | 10,950 |
| **Plasmacytoid DCs** | | | |
| Infection 6-day vs Ctrl | 1 | 1 | 11,612 |
| Infection 8-day vs Ctrl | 1 | 1 | 11,685 |
| **Plasma cells** | | | |
| Infection 6-day vs Ctrl | 3 | 0 | 8,844 |
| Infection 8-day vs Ctrl | 4 | 2 | 8,855 |
| **Stromal cells** | | | |
| Infection 6-day vs Ctrl | 0 | 0 | 10,499 |
| Infection 8-day vs Ctrl | 0 | 0 | 11,217 |
| **T cells** | | | |
| Infection 6-day vs Ctrl | 229 | 7 | 8,436 |
| Infection 8-day vs Ctrl | 28 | 128 | 6,237 |

MNPs, mononuclear phagocytes. DCs, dendritic cells.

**Table S2. Primers used in the construction of recombinant plasmids.**

| **Primer** | **Sequence (5' - 3')** |
| --- | --- |
| **Eukaryotic Igl-C3 expression** | |
| Igl-989-SgfI-S | CC GCGATCGC ATG GTTGAAGGCACTTCTACA |
| Igl-1088-Linker-AS | AGATCCTCCTCCGCC AATGCCTTTAGCTCCATT |
| FLAG-S | GGCGGAGGAGGATCT GATTACAAGGACGATGATGACA |
| FLAG-MluI-AS | CG ACGCGT TAA CTTGTCATCATCGTCCTTGTAATC |
| **Prokaryotic Igl-C3 expression** | |
| Igl-989-XhoI-S | CC CTCGAG GTTGAAGGCACTTCTACAGAA |
| Igl-1088-XhoI-AS | CC CTCGAG TTA AATGCCTTTAGCTCCATT |

The primers were synthesized by Invitrogen company.

**Table S3. Primers used in quantitative real-time RT-PCR (human).**

| **Species** | **Amplified gene** | | **Primer sequence (5' - 3')** |
| --- | --- | --- | --- |
| Human | *GAPDH* | S | TCACCACCATGGAGAAGGC |
|  |  | AS | GCTAAGCAGTTGGTGGTGCA |
| Human | *TNF* | S | AGCCCATGTTGTAGCAAACCC |
|  |  | AS | GAGGTACAGGCCCTCTGATG |
| Human | *IL1B* | S | CACGATGCACCTGTACGATCA |
|  |  | AS | GTTGCTCCATATCCTGTCCCT |
| Human | *IL6* | S | AGCCACTCACCTCTTCAGAAC |
|  |  | AS | GCCTCTTTGCTGCTTTCACAC |
| Human | *IL10* | S | GTGATGCCCCAAGCTGAGA |
|  |  | AS | CACGGCCTTGCTCTTGTTTT |
| Human | *IL23A* | S | ACACATGGATCTAAGAGAAGAGG |
|  |  | AS | CTATCAGGGAGCAGAGAAGG |
| Human | *TGFB1* | S | GTACCTGAACCCGTGTTGCT |
|  |  | AS | GTATCGCCAGGAATTGTTGC |
| Human | *CDH1* | S | GAGAACGCATTGCCACATACA |
|  |  | AS | ACCTTCCATGACAGACCCCTTAA |
| Human | *CLDN3* | S | CCACCAAGGTCGTCTACTC |
|  |  | AS | CCTGCGTCTGTCCCTTAGAC |
| Human | *TJP1* | S | TCGGCCAAATCTTCTCACTCC |
|  |  | AS | ACCAGTAAGTCGTCCTGATCC |
| Human | *MYD88* | S | CTAAGAAGGACCAGCAGAG |
|  |  | AS | GAAGCATCAGTAGGCATCA |
| Human | *LY96* | S | CACATTTTCTACATTCCAAG |
|  |  | AS | GTAATCGTCATCAGATCCTC |

The primers were synthesized by Invitrogen company.

**Table S3. Primers used in quantitative real-time RT-PCR (other species).**

| **Species** | **Amplified gene** | | **Primer sequence (5' - 3')** |
| --- | --- | --- | --- |
| Mouse | *Actb* | S | CACTGTCGAGTCGCGTCC |
|  |  | AS | TCATCCATGGCGAACTGGTG |
| Mouse | *Tnf* | S | GTCGTAGCAAACCACCAA |
|  |  | AS | GGCAGCCTTGTCCCTTGA |
| Mouse | *Il1b* | S | ACATCAGCACCTCACAAGCAG |
|  |  | AS | TTAGAAACAGTCCAGCCCATAC |
| Mouse | *Il6* | S | TGCCTTCTTGGGACTGAT |
|  |  | AS | TTGCCATTGCACAACTCTTT |
| Mouse | *Nos2* | S | TCCTGGAGGAAGTGGGCCGAAG |
|  |  | AS | CCTCCACGGGCCCGGTACTC |
| Mouse | *Arg1* | S | CAGAAGAATGGAAGAGTCAG |
|  |  | AS | CAGATATGCAGGGAGTCACC |
| *E. histolytica* | *Actin* | S | GCACTTGTTGTAGATAATGGATCAG |
|  |  | AS | ACCCATACCAGCCATAACTGAAACG |
| *E. histolytica* | *Igl1* | S | TCTTGTAATAAGTTCCCGGAGCA |
|  |  | AS | CATCAGAAACAGTACATCTTTTATTACATG |
| *E. histolytica* | *Igl2* | S | GTACTAAATACCCAGATCATTGTTCAAA |
|  |  | AS | CATCAGAAACAGTACATCTTTTATTACATG |

The primers were synthesized by Invitrogen company.

**Table S4. Antibodies used in western blotting.**

| **Reagent** | **Source** | **Identifier** |
| --- | --- | --- |
| **Primary antibodies** | | |
| β-Actin (8H10D10) Mouse mAb | Cell Signaling Technology | #3700 |
| TNF-α Rabbit Polyclonal antibody | Cell Signaling Technology | #3707 |
| IL-1β (D3U3E) Rabbit mAb | Cell Signaling Technology | #12703 |
| IL-6 (D3K2N) Rabbit mAb | Cell Signaling Technology | #12153 |
| MyD88 (D80F5) Rabbit mAb | Cell Signaling Technology | #4283 |
| NF-κB p65 (D14E12) XP® Rabbit mAb | Cell Signaling Technology | #8242 |
| Phospho-NF-κB p65 (Ser536) (93H1) Rabbit mAb | Cell Signaling Technology | #3033 |
| LY96/MD2 Rabbit Polyclonal antibody | Proteintech | 11784-1-AP |
| *E. histolytica* Igl-immunized hamster serum | Lab-prepared | NA |
| **Secondary antibodies** | | |
| Goat Anti-Mouse IgG H&L (HRP) | Abcam | Ab6789 |
| Goat Anti-Rabbit IgG H&L (HRP) | Abcam | Ab6721 |
| Goat Anti-Syrian Hamster IgG H&L (HRP) | Abcam | Ab6892 |

**Table S5. Oligonucleotides used in small interfering RNA treatment.**

| **Oligonucleotide** | | **Sequence (5' - 3')** |
| --- | --- | --- |
| Control siRNA | S | UUCUCCGAACGUGUCACGUTT |
|  | AS | ACGUGACACGUUCGGAGAATT |
| Human *GAPDH* siRNA | S | GUAUGACAACAGCCUCAAGTT |
|  | AS | CUUGAGGCUGUUGUCAUACTT |
| Human *MYD88* siRNA | S | GGCAACUGGAACAGACAAATT |
|  | AS | UUUGUCUGUUCCAGUUGCCTT |
| Human *LY96* siRNA | S | GACUGUGAAUACAACAAUATT |
|  | AS | UAUUGUUGUAUUCACAGUCTT |

The oligonucleotides were synthesized by Invitrogen company.
