## Supplementary material for "*Entamoeba histolytica* Gal/GalNAc lectin intermediate subunit promotes inflammation and epithelial damage in intestinal amebiasis through its C3 region": Figure Legends

**Fig. 1** Intense inflammatory response in the C3H/HeNCrl murine model of intestinal amebiasis. (a) Schematic of experimental design for the murine intestinal amebiasis model. (b) Weight change over the experimental period in amoeba-infected mice (red), exhibiting the weight ratio of each date to Day 0. Control-treated mice (black) were inoculated with the same volume of amoebic culture medium. (c) The proportions of neutrophil (NEUT), monocyte (MONO), lymphocyte (LYMPH), and eosinophil (EO) to all leukocytes in the peripheral blood of different amoeba-infected and control-treated mice. (d) Representative immunohistochemical images of TNF-α, IL-1β, and IL-6 staining and their quantification in the mouse ceca in each group (n = 3). Scale bar: 25 μm. (e) Cell type assignment of all 68,058 single cells from the ceca of nine different mice (3 animals in each of 3 groups) visualized using a UMAP plot. (f) Number of DEGs per cell type in the ceca of amoeba-infected and control-treated mice. Upregulated DEGs (log2FC value > 0.585 and adjusted p value < 0.05) are displayed in red, downregulated DEGs (log2FC value < -0.585 and adjusted p value < 0.05) are displayed in blue, and other genes (|log2FC| value > 0.585 and adjusted p value ≥ 0.05) are displayed in grey. (g) GSEA of GO terms identifying broad enrichment of gene sets associated with inflammation in MNPs. Filter criteria: |NES| > 1, adjusted p value < 0.05, FDR < 0.25. (h) GSEA of GO terms related to energy metabolism, inflammation, and cell-cell junction in epithelial cells. Filter criteria: |NES| > 1, adjusted p value < 0.05, FDR < 0.25. Data are expressed as mean with SD. *p < 0.05; **p < 0.01; ***p < 0.001; ****p < 0.0001.

**Fig. 2** Mononuclear phagocytes play an important role in intestinal amebiasis. (a) UMAP visualization of all intestinal single cells, with the Amebiasis pathway (mmu05146) in KEGG enrichment analysis being scored and color-coded (quantile normalized). (b) Violin plot displaying AUCell scores of the KEGG Amebiasis pathway gene set for all cell types in each group. Quantile normalization was performed to assess the gene set activity. (c) Ratio of cell type percentages in each group. (d) Top five significantly upregulated KEGG pathways in GSEA comparing Infection 6-day group and Ctrl group MNPs (sorted by NES). Changes of the core enriched genes for GSEA enriched categories are shown in the ridgeline plot. (e) GSEA showing the gene enrichment of the KEGG Amebiasis pathway compared between Infection 6-day group and Ctrl group MNPs and the histogram of changes in its core enriched genes. (f) Ridgeline plot of core enrichment gene changes of the top five significantly upregulated KEGG pathways in GSEA compared between Infection 8-day group and Ctrl group MNPs (sorted by NES). (g) GSEA showing the gene enrichment of the KEGG Amebiasis pathway compared between Infection 8-day group and Ctrl group MNPs. Changes in the corresponding core enriched genes are exhibited in the histogram. (h) qRT-PCR assay of *TNF*, *IL1B*, and *IL6* expressions in human U937 cells preincubated with *E. histolytica* trophozoites for 60 min. (i) qRT-PCR assay of *TNF*, *IL1B*, and *IL6* expressions in human U937 cells preincubated with native Igl protein for 24 h. Data are expressed as mean with SD.

**Fig. 3** The C3 segment of *E. histolytica* Igl induces an inflammatory response in macrophages. (a) qRT-PCR assay of *TNF*, *IL1B*, *IL6*, *IL10*, *IL23A*, and *TGFB1* expressions in human U937 cells preincubated with eukaryotic Igl-C3 (2 μg/mL), LPS (100 ng/mL), or IL-4 (20 ng/mL) for 24 h. (b) Immunoblots of TNF-α, IL-1β, and IL-6 in control and different Igl-C3-treated U937 cells. β-Actin serves as the loading control. (c) Quantification of TNF-α, IL-1β, IL-6, IL-10, IL-23, and TGF-β1 concentrations in U937 cell culture supernatant by corresponding ELISA kits following preincubation with eukaryotic Igl-C3. (d) Principal component analysis of the transcriptomic sequencing data of Control 12 h group, Control 24 h group, Igl-C3-treated 12 h group, and Igl-C3-treated 24 h group human U937 cells (n = 3). (e) Volcano plot of DEGs in Igl-C3-treated 12 h group versus Control 12 h group U937 cells (|log2FC| value > 1 and adjusted p value < 0.05). (f) Volcano plot of DEGs in Igl-C3-treated 24 h group versus Control 24 h group U937 cells (|log2FC| value > 1 and adjusted p value < 0.05). (g) Heatmap exhibiting the changes of cytokine gene expressions in U937 cells after 12 h and 24 h incubation with eukaryotic Igl-C3. Data are expressed as mean with SD. *p < 0.05; **p < 0.01; ***p < 0.001; ****p < 0.0001.

**Fig. 4** Indirect damage of Igl-C3 to intestinal epithelium integrity by inducing macrophage inflammation. (a) Circle plots showing the number of interactions between MNPs and other cell types in each group of the single-cell RNA-seq based on the CellChat database. With MNPs as the sender and other cell types as the receivers, linewidth corresponds to the number of interactions. (b) Circle plot exhibiting the differential numbers of interactions between various cell types in Infection 6-day group versus Ctrl group or Infection 8-day group versus Ctrl group comparison. Red lines represent upregulated cell-cell interactions and blue lines represent downregulated cell-cell interactions, while linewidth corresponds to the overall change in interaction numbers. (c) Detailed changes of molecular interactions between MNPs and epithelial cells in Infection 6-day group versus Ctrl group or Infection 8-day group versus Ctrl group comparison. In all interactions, the MNP population is on the y-axis, and epithelial cell population is on the x-axis. (d) GSEA showing the gene enrichment of Tight Junction term (GO:0070160) in GO enrichment analysis compared between Infection 6-day group and Ctrl group or between Infection 8-day group and Ctrl group epithelial cells. (e) Histogram of changes in gene expressions of the *Cdh*, *Cldn*, and *Tjp* gene family members compared between Infection 6-day group and Ctrl group or between Infection 8-day group and Ctrl group epithelial cells. Filter criteria: |log2FC| value > 0.585, adjusted p value < 0.05. (f) Brief illustration for the establishment of three-dimensional triple culture system. The system consists of an epithelial cell monolayer (Caco-2 and HT29-MTX-E12 cells) on top of the transwell inserts and macrophages (U937 cells) in culture plate wells to mimic the structure of intestinal epithelium and lamina propria macrophages. (g) Observation by light microscopy of the intestinal epithelial model stained with alcian blue and eosin. Alcian blue was used to visualize acidic mucins. (h) Transepithelial electronic resistance measurement of Igl-C3 (2 μg/mL)-treated epithelial cell monolayers in the presence or absence of macrophages. (i) Permeability to 4 kDa FITC-Dextran of Igl-C3-treated epithelial cell monolayers in the presence or absence of macrophages. (j) qRT-PCR assay of *CDH1*, *CLDN3*, and *TJP1* expressions in Caco-2 or HT29-MTX-E12 cells preincubated with Igl-C3 protein. U937 cells were added as mentioned above. Data are expressed as mean with SD. *p < 0.05; **p < 0.01; ****p < 0.0001.

**Fig. 5** Macrophages recognize Igl-C3 via TLR4 and initiate the TLR4/MyD88/NF-κB inflammatory signaling pathway. (a) Interaction network between the top 20 pathways with the most significant upregulation (sorted by adjusted p value) in KEGG enrichment analysis. Igl-C3-treated 12 h group and Control 12 h group in U937 cell transcriptomic sequencing were compared. (b) Interaction network between the top 20 KEGG pathways with the most significant upregulation in Igl-C3-treated 24 h group versus Control 24 h group U937 cells. (c) qRT-PCR assay of *TNF*, *IL1B*, and *IL6* expressions in human U937 cells treated with Igl-C3 (2 μg/mL) for 24 h. Cells were preincubated with TLR4 inhibitor (TAK-242, 1 μM) or TLR1/2 inhibitor (CU-CPT22, 1 μM) for 2 h before Igl-C3 treatment. (d) Immunoblots of MyD88, NF-κB, and phospho-NF-κB in control and different Igl-C3-treated U937 cells. β-Actin serves as the loading control. (e) Observation by laser confocal microscopy of phospho-NF-κB content and distribution in U937 cells preincubated with Igl-C3. Cell nuclei were counterstained with DAPI. (f) qRT-PCR validation of *MYD88* and *GAPDH* expressions in human U937 cells after different siRNA transfections (40 nM). (g) Immunoblots of MyD88, NF-κB, and phospho-NF-κB in Igl-C3-treated U937 cells after different siRNA transfections. β-Actin serves as the loading control. (h) Observation by laser confocal microscopy of phospho-NF-κB content and distribution in Igl-C3-treated U937 cells after different siRNA transfections. Cell nuclei were counterstained with DAPI. (i) qRT-PCR assay of *TNF*, *IL1B*, and *IL6* expressions in Igl-C3-treated U937 cells after different siRNA transfections. Data are expressed as mean with SD. *p < 0.05; **p < 0.01; ***p < 0.001; ****p < 0.0001.

**Fig. 6** Igl-C3 binds to the TLR4 co-receptor MD2 on the surface of macrophages to activate inflammatory signals. (a) qRT-PCR assay of *TNF*, *IL1B*, and *IL6* expressions in human U937 cells treated with Igl-C3 (2 μg/mL) for 24 h. Cells were preincubated with MD2 inhibitor (MD2-IN-1, 5 μM) for 2 h before Igl-C3 treatment. (b) Immunoblots of MyD88, NF-κB, phospho-NF-κB, TNF-α, IL-1β, and IL-6 in control and Igl-C3-treated U937 cells after preincubation with the MD2 inhibitor. β-Actin serves as the loading control. (c) Quantification of TNF-α, IL-1β, IL-6, IL-10, IL-23, and TGF-β1 concentrations in U937 cell culture supernatant by corresponding ELISA kits following preincubation with the MD2 inhibitor and treatment with Igl-C3 protein. (d) qRT-PCR validation of *LY96* (MD2) and *GAPDH* expressions in human U937 cells after different siRNA transfections (40 nM). (e) Western blotting validation of MD2 expression in human U937 cells after different siRNA transfections. β-Actin serves as the loading control. (f) qRT-PCR assay of *TNF*, *IL1B*, and *IL6* expressions in Igl-C3-treated U937 cells after different siRNA transfections. (g) Affinities of human MD2 protein to eukaryotic Igl-C3, prokaryotic Igl-C3, and LPS analyzed by ELISA. (h) Biacore analysis for detecting the binding ability of eukaryotic Igl-C3 to human MD2 and TLR4 proteins. (i) Biacore analysis for detecting the binding ability of native Igl to human MD2 protein. Data are expressed as mean with SD. *p < 0.05; **p < 0.01; ***p < 0.001; ****p < 0.0001.

**Fig. 7** *E. histolytica* Igl diffuses along with EVs of the parasite to stimulate host macrophages. (a) Coomassie Brilliant Blue staining of amoebic EV samples and corresponding non-reduced and reduced immunoblots to confirm the presence of Igl protein. (b) Venn diagram of protein compartments and histogram of protein quantities of amoebic EVs in macrophage coculture group trophozoites and control culture group trophozoites. Quantitative proteomics analysis was performed on EV protein samples of the two groups. (c) Box diagram showing the fraction of total (FOT) protein abundance in quantitative results. Horizontal line in the middle of each box represents the median value. (d) Heatmap exhibiting correlations between any two samples in each group. (e) Volcano plot of differential proteins in macrophage coculture group versus control culture group amoebic EVs. Filter criteria: |log2FC| value > 1, adjusted p value < 0.05. (f) Upregulated changes of terms under Molecular Function in the GO enrichment analysis compared between macrophage coculture group and control culture group amoebic EVs. (g) Downregulated changes of the GO terms under Molecular Function in macrophage coculture group versus control culture group amoebic EVs. (h) Interaction network of upregulated differential proteins in amoebic EVs after coculture with host macrophages based on the STRING database. (i) Changes of Gal/GalNAc lectin component proteins in amoebic EVs after coculture with host macrophages. (j) qRT-PCR assay of *Igl1* and *Igl2* expressions in *E. histolytica* trophozoites after 60 min coculture with host macrophages. Data are expressed as mean with SD. *p < 0.05.

**Fig. 8** Graphical summary of the underlying mechanism found in this study. In addition to detachment from the membrane surface and trophozoite rupture, *E. histolytica* Igl can also diffuses along with amoebic EVs, forming the functional basis of its pathogenic effects. On the surface of macrophages, Igl binds to the TLR4 co-receptor MD2 through its C3 region, initiating the TLR4/MyD88/NF-κB inflammatory signaling pathway in host cells. By inducing macrophage inflammation and cytokine production, *E. histolytica* indirectly impairs intestinal epithelium integrity through the Igl protein. In response to the parasite invasion during intestinal amebiasis, the present study provides new insights elucidating the inflammatory formation and epithelial damage of gut mucosal immune system.

**Fig. S1** Quality control and cell type annotation of single-cell RNA-seq data. (a) Number of total expressed genes (nFeature_RNA), number of transcripts (nCount_RNA), percentage of mitochondria genes (percent.mito), percentage of ribosomal genes (percent.ribo), and percentage of red blood cell genes (percent.Hb) of individual cells analyzed to filter out low-quality information after single-cell sequencing. (b) Clustering of single cells in the mouse cecum visualized using a UMAP plot. (c) Proportion of the 51 cell clusters in each group (quantile normalized). (d) Marker genes for cell type annotation in all clusters. Dot size represents percentage of cells expressing the particular gene within the cell type, while dot color represents average expression level. (e) H&E staining of mouse cecal tissue observed by light microscopy. The area of cecal lymph node in the image was further zoomed in. Scale bar: 500 μm.

**Fig. S2** KEGG enrichment analysis of mononuclear phagocytes in single-cell RNA-seq data. (a) All significantly upregulated KEGG pathways in GSEA comparing Infection 6-day group and Ctrl group MNPs (sorted by NES). (b) All significantly upregulated KEGG pathways in GSEA comparing Infection 8-day group and Ctrl group MNPs (sorted by NES). (c) All significantly downregulated KEGG pathways in GSEA comparing Infection 6-day group and Ctrl group MNPs (sorted by NES). (d) All significantly downregulated KEGG pathways in GSEA comparing Infection 8-day group and Ctrl group MNPs (sorted by NES).

**Fig. S3** Igl-C3 induces an inflammatory response in different macrophage cell lines. (a) qRT-PCR assay of *TNF*, *IL1B*, *IL6*, *IL10*, *IL23A*, and *TGFB1* expressions in PMA-differentiated human THP-1 cells preincubated with eukaryotic Igl-C3 (2 μg/mL), LPS (100 ng/mL), or IL-4 (20 ng/mL) for 24 h. (b) qRT-PCR assay of *Tnf*, *Il1b*, *Il6*, *Nos2*, and *Arg1* expressions in mouse RAW264.7 cells preincubated with eukaryotic Igl-C3 (2 μg/mL) for 24 h. Data are expressed as mean with SD. *p < 0.05; **p < 0.01; ****p < 0.0001.

**Fig. S4** Quality control and GO enrichment analysis of transcriptomic sequencing data of different Igl-C3-treated U937 cells. (a) Box diagram of the gene expression level in each sample using StringTie software version 1.3.1c. The first quartile, median, and third quartile values are shown. (b) Heatmap exhibiting correlations between any two samples from four groups. (c) Top 10 significantly altered GO terms under each category and their rich factors comparing Igl-C3-treated 12 h group and Control 12 h group U937 cells (sorted by adjusted p value). (d) Top 10 significantly altered GO terms under each category and their rich factors comparing Igl-C3-treated 24 h group and Control 24 h group U937 cells (sorted by adjusted p value).

**Fig. S5** KEGG enrichment analysis of transcriptomic sequencing data of different Igl-C3-treated U937 cells. (a) Top 20 significantly altered KEGG pathways and their rich factors comparing Igl-C3-treated 12 h group and Control 12 h group U937 cells (sorted by adjusted p value). Percentage of the DEG number belonging to each KEGG pathway to all candidate DEGs was exhibited (|log2FC| value > 1 and adjusted p value < 0.05). (b) Histogram of DEG changes in the KEGG Amebiasis pathway (hsa05146) comparing Igl-C3-treated 12 h group and Control 12 h group U937 cells. (c) Top 20 significantly altered KEGG pathways and their rich factors comparing Igl-C3-treated 24 h group and Control 24 h group U937 cells (sorted by adjusted p value). Percentage of the DEG number belonging to each KEGG pathway was shown as described above. (d) Histogram of DEG changes in the KEGG Amebiasis pathway comparing Igl-C3-treated 24 h group and Control 24 h group U937 cells.

**Fig. S6** Signal sending and receiving for each cell type in single-cell RNA-seq data based on the CellChat database. (a) Incoming signaling patterns of cellular communications among various cell types in each group. (b) Outgoing signaling patterns of cellular communications among various cell types in each group.

**Fig. S7** Immunohistochemical staining of tight junction-associated proteins in the mouse ceca. (a) Representative images of E-cadherin staining and the quantification in each group. (b) Representative images of Claudin-3 staining and the quantification in each group. (c) Representative images of ZO-1 staining and the quantification in each group. Scale bar: 50 μm. Data are expressed as mean with SD. **p < 0.01; ***p < 0.001; ****p < 0.0001.

**Fig. S8** Gene set enrichment analysis of the KEGG Toll-like receptor signaling pathway (mmu04620) compared between different groups of mononuclear phagocytes in single-cell RNA-seq data. (a) GSEA showing the gene enrichment of KEGG Toll-like receptor signaling pathway compared between Infection 6-day group and Ctrl group MNPs and the histogram of changes in its core enriched genes. (b) GSEA showing the gene enrichment of KEGG Toll-like receptor signaling pathway compared between Infection 8-day group and Ctrl group MNPs and the histogram of changes in its core enriched genes.

**Fig. S9** Protein abundance of amoebic EVs in different groups and the interaction network of differential proteins in quantitative proteomics. (a) Top 20 most abundant proteins found in control culture group amoebic EVs. (b) Top 20 most abundant proteins found in macrophage coculture group amoebic EVs. (c) Interaction network of downregulated differential proteins in amoebic EVs after coculture with host macrophages based on the STRING database.
